# Persistent DNA Repair Signaling and DNA Polymerase Theta Promote Broken Chromosome Segregation

**DOI:** 10.1101/2021.06.18.449048

**Authors:** Delisa E. Clay, Heidi S. Bretscher, Erin A. Jezuit, Korie B. Bush, Donald T. Fox

**Author notes:** email for correspondence, Mailing address: C318 LSRC, DUMC Box 3813, Duke University Medical Center, Durham NC 27710, USA.

## Abstract

Cycling cells must respond to double-strand breaks (DSBs) to avoid genome instability. Mis-segregation of chromosomes with DSBs during mitosis results in micronuclei, aberrant structures linked to disease. How cells respond to DSBs during mitosis is incompletely understood. We previously showed that *Drosophila* papillar cells lack DSB checkpoints (as observed in many cancer cells). Here, we show that papillar cells still recruit early-acting repair machinery (Mre11 and RPA3) to DSBs. This machinery persists as foci on DSBs as cells enter mitosis. Repair foci are resolved in a step-wise manner during mitosis. Repair signaling kinetics at DSBs depends on both monoubiquitination of the Fanconi Anemia (FA) protein Fancd2 and the alternative end-joining protein DNA Polymerase Theta. Disruption of either or both of these factors causes micronuclei after DNA damage, which disrupts intestinal organogenesis. This study reveals a mechanism for how cells with inactive DSB checkpoints can respond to DNA damage that persists into mitosis.

**Summary:** Clay et. al. show that cells with DNA breaks that persist into mitosis activate sustained DNA repair signaling, regulated by Fanconi Anemia proteins and the alternative end-joining repair protein DNA Polymerase Theta. This signaling enables broken chromosome segregation and prevents micronuclei.

## Introduction

Cells are constantly at risk for DNA damage from internal and external factors (Cannan and Pederson, 2016), and must respond to such damage to maintain genome stability (Ciccia and Elledge, 2010; Harper and Elledge, 2007). DNA lesions trigger DNA damage responses (DDRs) that activate checkpoints that result in cell cycle arrest, DNA repair, or apoptosis (Jackson and Bartek, 2009; Petsalaki and Zachos, 2020). These DDR checkpoints primarily act in interphase but allow for faithful chromosome segregation when damaged cells resume cycling and enter mitosis (Finn et al., 2012; Sekelsky, 2017). If cell cycle checkpoints are inactivated or dysregulated, this can pose a threat to genome integrity in that damaged DNA can persist into mitosis (Hafner et al., 2019; Santivasi and Xia, 2014). One of the more problematic forms of persistent DNA damage for the mitotic cell are double-strand breaks (DSBs), which can lead to mis-segregation and subsequent loss of portions of the genome (Aleksandrov et al., 2020; Vignard et al., 2013). DSBs generate acentric DNA, a DNA fragment that lacks canonical kinetochore-spindle attachments and is therefore at an increased risk of mis-segregating if present during mitosis (Kanda et al., 1998; Warecki and Sullivan, 2020). Failure to properly segregate acentric DNA during mitosis can lead to micronuclei, aberrant nuclear structures characterized by a poorly-formed nuclear envelope (Bretscher and Fox, 2016; Crasta et al., 2012; Durante and Formenti, 2018).

However, acentric DNA can segregate properly in several organisms including yeast, *Drosophila,* and human cells (Warecki and Sullivan, 2020). In *S. pombe,* acentric DNA can segregate in mitosis by forming a neocentromere and by noncanonical DNA repair (Ishii et al., 2008; Ohno et al., 2015). In *C. elegans,* acentric meiotic chromosomes are thought to segregate through microtubule-generated forces driving poleward movement (Dumont et al., 2010). In *Drosophila* brain progenitor cells, acentric fragments can segregate poleward during mitosis using various mechanisms involving protein-based tethers, microtubule generated forces, nuclear envelope reformation and fusion, as well as the recruitment of early-acting repair proteins (Derive et al., 2015; Karg et al., 2017; Karg et al., 2015; Landmann et al., 2020; Royou et al., 2010; Warecki et al., 2020). Therefore, there are several proposed mechanisms for how acentric DNA can properly segregate during mitosis under distinct conditions.

Despite the various mechanisms identified for how acentric DNA segregates properly during mitosis and avoids micronuclei formation, questions still remain surrounding possible physical connections that might link the acentric fragment to segregating, centromeric DNA. More specifically, DNA repair signaling could potentially regulate such a linkage, or influence segregation behavior of acentric DNA. We established *Drosophila* hindgut rectal papillar cells (hereafter: papillar cells) as an accessible model to understand cellular responses to damaged, acentric DNA that is present during mitosis (Bretscher and Fox, 2016). We previously found that papillar cells inactivate checkpoint responses to DNA damage at a specific developmental time point. At the second larval instar stage (L2), papillar cells undergo two rounds of endocycling (Fox and Duronio, 2013; Øvrebø and Edgar, 2018), where they replicate their genome content without cell division (**Fig. 1A, L2**). During this stage, papillar cells do not arrest the cell cycle or undergo apoptosis in response to high levels of DNA damage (20 Gy X-ray Irradiation, IR), as in other endocycling cells (Calvi, 2013; Hassel et al., 2014; Mehrotra et al., 2008). Papillar cells do not respond to changes in p53 expression nor do they depend on the checkpoint kinases Chk1 and Chk2 following DNA damage (Bretscher and Fox, 2016). Despite inactivating canonical DNA damage responses, several (5-6) days later during the early pupation stage, papillar cells leave a G2-like state and enter into a mitotic cell cycle (**Fig. 1A, Early pupal,** (Fox et al., 2010; Stormo and Fox, 2016; Stormo and Fox, 2019), As a consequence, papillar cells frequently enter mitosis with acentric DNA fragments (∼12% of divisions; Bretscher and Fox, 2016). Despite papillar cell acentric DNA lacking obvious physical connections, these fragments lag but ultimately segregate into daughter nuclei during mitosis (**Fig 1A, mitotic**). We identified that papillar cell acentric DNA segregation requires proteins from the Fanconi Anemia (FA) pathway, Fancd2 and FancI (Bretscher and Fox, 2016).

**Figure 1.**
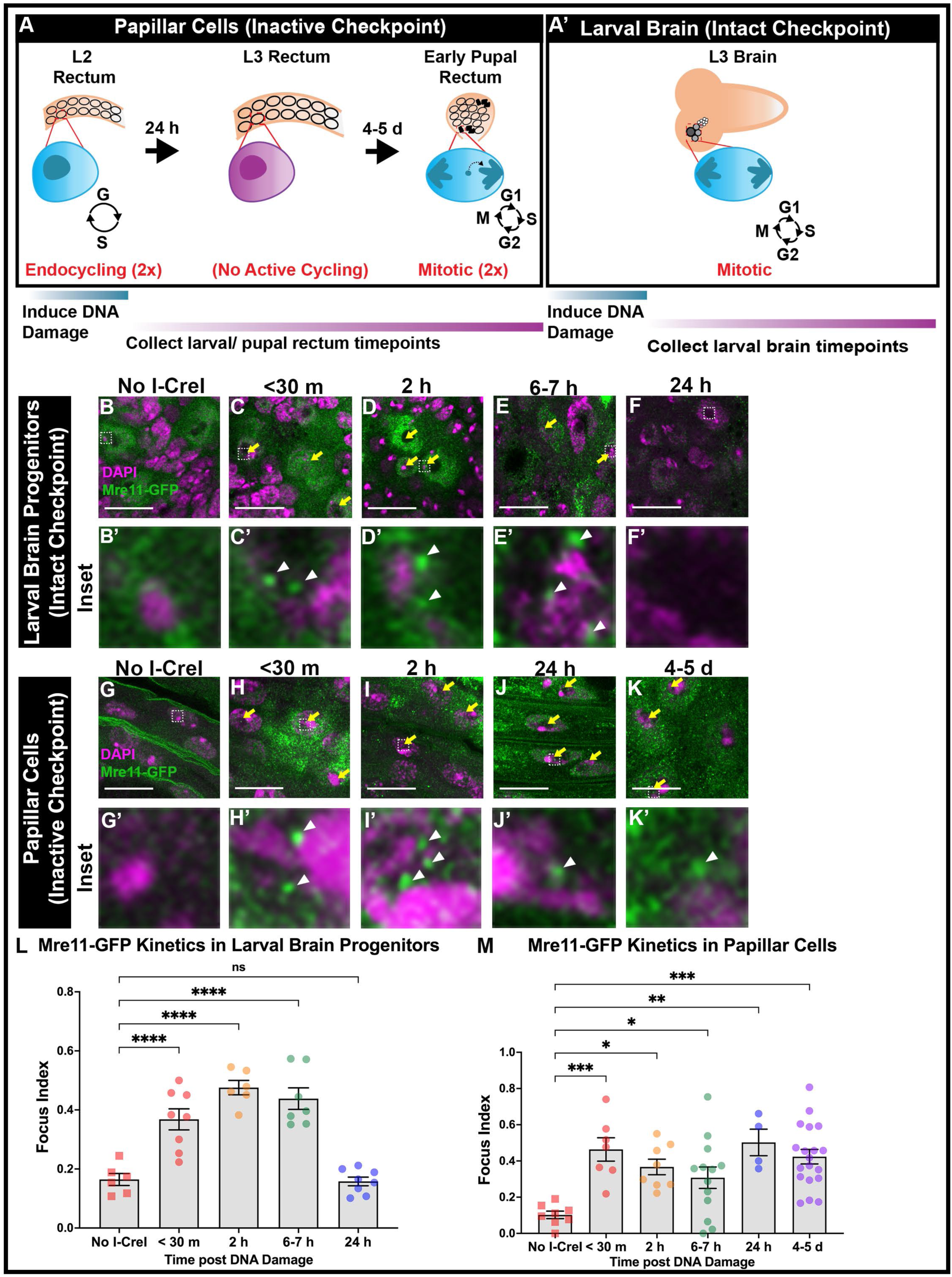
Mre11 is recruited to checkpoint inactivated papillar cells and persists for days. (**A**) Schematic of the developing *Drosophila* rectum from the second larval instar (L2) stage to early pupation. DNA damage is induced during L2 stage when checkpoints are inactive. Various timepoints are collected until mitosis 4-5 d post L2. (**A’**) Schematic of the feeding third larval instar (L3) brain. DNA damage is induced during the L3 stage. Larval brain progenitors (neuroblasts [dark gray] and ganglion mother cells [gray]) are actively cycling and have intact checkpoints. Various timepoints are collected for 24 h. (**B-F’**) Mre11 (Mre11-GFP) localization over time +/− *hs*-*I-Cre*I in larval brain progenitors. Time after break induction is indicated in minutes (m) or hours (h). Mre11+ foci are marked with yellow arrows. The hatched box highlights an area magnified 10X in the corresponding inset below each panel. Enlarged foci are marked with white arrowheads. (**G-K’**) Mre11-GFP localization over time in checkpoint-inactive papillar cells +/− *hs*-*I-Cre*I. Labeling as in **B-F’.** Scale bars = 10μm. (**L,M**) Quantification of Mre11+ foci kinetics in larval brain stem cells (**L**) and papillar cells (**M**). Focus index is the frequency of cells with at least one focus. Each data point represents one animal. Each timepoint has at least 2 replicates. Statistical test: Ordinary one-way ANOVA, p-value for (**L**) <0.0001. P-value for (**M**) p=0.0002. See Methods for statistical notations. All images in figure: Mre11, green; DNA (DAPI), magenta.

In this current study, we further investigate the mechanism of acentric DNA segregation to better understand possible linkage of acentric fragments to centromeric DNA during mitosis. We find that despite lacking a canonical DDR, papillar cells recruit early-acting repair proteins from the MRN (Mre11) and RPA (RPA3) complexes to DSBs. These proteins persist on damaged papillar chromosomes for days (5-6 days) following induced DSBs and remain present as these cells enter mitosis. During mitosis, Mre11 and RPA3 show distinct kinetics, with Mre11 leaving the DNA prior to RPA3. Further, we find that the alternative end-joining repair protein DNA Polymerase Theta (hereafter Pol Theta, encoded by the *polQ* gene), but not homologous recombination nor canonical homologous end-joining, is required for RPA3 removal during mitosis, for acentric DNA segregation, and for micronuclei prevention. Finally, we show that *polQ* is epistatic to the conserved monoubiquitination of Fancd2. The action of Mre11, Pol Theta, and monoubiquitinated Fancd2 are critical for intestinal organogenesis following DSBs. Our findings highlight a role for persistent DNA repair signaling, regulated by a conserved alternative end-joining protein and monoubiquitinated Fancd2, on acentric DNA.

## Results

### Mre11 and RPA3 are recruited to damaged papillar chromosomes but are not resolved prior to mitosis

Although papillar cells lack apoptotic and cell cycle arrest responses to DNA damage, it remained possible that these cells activate DNA repair responses to segregate acentric DNA. To visualize DNA repair signaling dynamics, we investigated the temporal recruitment of early-acting DNA damage repair proteins following double strand DNA breaks (DSBs). The highly conserved MRN (Mre11, Rad50, Nbs) complex is recruited immediately following a DSB and is required for initiating downstream repair events (Petsalaki and Zachos, 2020; Syed and Tainer, 2018; Tisi et al., 2020). Using animals expressing ubi-*mre11-GFP* (Landmann et al., 2020), we assayed the localization of this MRN component over time to papillar chromosomes with DSBs. As performed in our previous studies and by other groups (Royou et al., 2010; Royou et al., 2005), we induced targeted DSBs using the endonuclease, *hs*-*I-Cre*I. *hs*-*I-Cre*I creates DSBs in the rDNA of the *Drosophila* sex chromosomes (Rong et al., 2002; Royou et al., 2010). Using *hs*-*I-Cre*I, we are able to induce DSBs in papillar cell chromosomes, increasing the frequency of cells with acentric DNA from 12% to up to 90% (Bretscher and Fox, 2016).

As a control for mitotic cycling cells with an intact DSB response, we examined third larval instar (L3) stage brain progenitor cells (neuroblast and ganglion mother cells, **Fig. 1A’,** Jaklevic et al., 2006; Peterson et al., 2002; Royou et al., 2005). In larval brain progenitors, Mre11+ foci are recruited shortly (<30 minutes, m) following DSB induction using *hs*-*I-Cre*I (**Fig. 1 B, B’ vs. C,C’, L**). These foci are mostly resolved 24 hours (h) after DNA damage, suggesting that repair is complete within this timeframe (**Fig. 1C-F’,L**). Consistent with this idea, 24 h after DNA damage, we observe mitotic neural progenitors, suggesting that these cells begin to exit cell cycle arrest (data not shown). Similar to our findings in the brain, checkpoint-inactive papillar cells also recruit Mre11+ foci within 30 m after inducing DSBs using *hs*-*I-Cre*I (**Fig. 1 G, G’ vs. H, H’, M**). However, these foci persist for much longer than in larval brain progenitors (**Fig. 1H-K,M**). During the early pupal stage, several days after we induce DSBs in papillar cells, Mre11+ foci still persist on damaged papillar chromosomes. The long time-frame of Mre11 persistence corresponds with the 5-6 day (d) period between papillar cell endocycles and the first mitotic division, which occurs 24 h after pupation onset (**Fig. 1A**, Bretscher and Fox, 2016). Thus, although an MRN component is promptly recruited to checkpoint-inactive papillar cells shortly after a DSB, this protein persists on broken DNA into mitosis.

We next investigated whether Mre11 recruitment coincides with and recruits other markers of DSB repair. Mre11+ foci in both brain and papillar cells colocalizes with !H2Av, a hallmark of DSB signaling (**Fig. S1A-B’; S1D-E’**). The MRN complex is known to bind to either side of a DSB, holding the broken ends in close proximity. MRN can then recruit various downstream repair factors leading to several different DSB repair pathways. MRN can also initiate DNA resection, a process in which 5’ DNA ends are processed leaving 3’ single-strand DNA overhangs. This in turn can generate DNA templates necessary for multiple DSB repair pathways. These 3’ overhangs are then coated by the heterotrimeric complex RPA (RPA1, RPA2, RPA3), which prevents annealing of single-strand DNA (Syed and Tainer, 2018; Tisi et al., 2020).

To determine if MRN recruits RPA in papillar cells after DNA damage, we used animals expressing *RPA3-GFP* with a ubiquitous promoter (*ubi-RPA3-GFP*, Murcia et al., 2019). We find that similar to Mre11, RPA3+ foci are observed in less than 30 m after DNA damage is induced using *hs*-*I-Cre*I in checkpoint-intact larval brain progenitors (**Fig. 2 A, A’ vs. B,B’,D**). These foci are then resolved 24 h after inducing DNA damage (**Fig. 2C, C’, D**). RPA3+ foci also appear in papillar cells within 30 m following *hs*-*I-Cre*I induced DNA damage (**Fig. 2 E, E’ vs. F, F’, J**). However, unlike in neural progenitors, these foci are not resolved by 24 h (**Fig. 2G, G’, J**) and *hs*-*I-Cre*I induced foci persist for 4-5 d after DNA damage (**Fig. 2H, H’, J)** similar to Mre11 kinetics in papillar cells.

**Figure 2.**
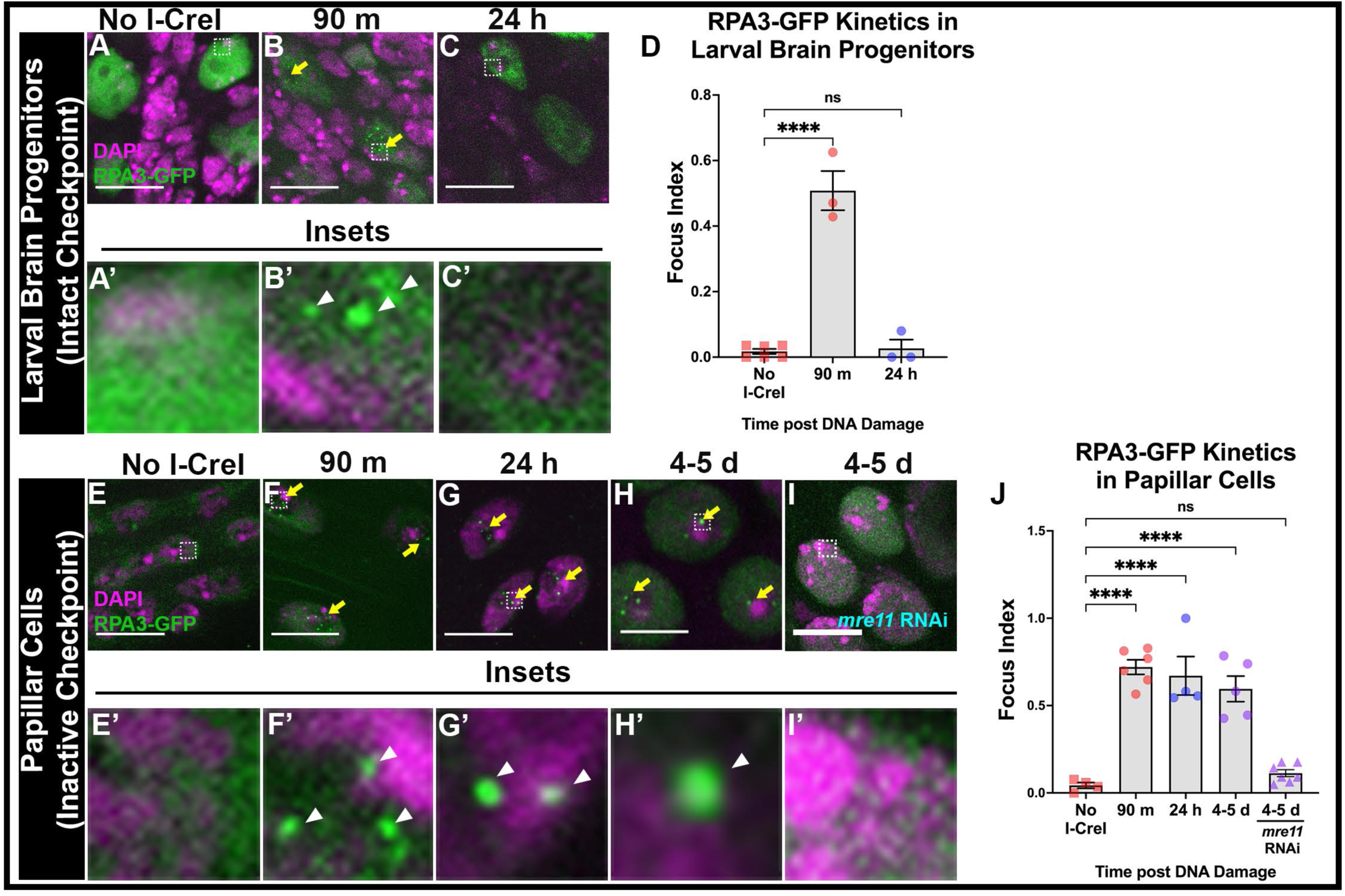
RPA3 is recruited to checkpoint inactive papillar cells and persists for days. (**A-C’**) RPA3 localization over time in larval brain progenitors +/− *hs*-*I-Cre*I. Time after break induction is indicated in minutes (m) or hours (h). RPA3+ foci are marked with yellow arrows. The hatched box highlights an area magnified 10X in the corresponding inset below each panel. Enlarged foci are marked with white arrowheads. Scale bars = 10μm. (**D**) Quantification of RPA3+ foci kinetics in larval brain progenitors. (**E-I’**) RPA3-GFP localization over time in WT papillar cells (**E-H’**) *mre11* RNAi expressing papillar cells (**I,I’**) and +/− *hs*-*I-Cre*I. Labeling as in **A-C’**. (**J**) Quantification of RPA3+ foci kinetics in papillar cells. Each data point represents one animal. Each timepoint has at least 2 replicates. Statistical test: Ordinary one-way ANOVA, p-value for (**D**) p<0.0001. P-value for (**J**) p<0.0001. See Methods for statistical notations. All images in figure: RPA3, green; DNA (DAPI), magenta.

We then determined if RPA3 is recruited to papillar DSBs in an Mre11-dependent manner. We examined RPA3-GFP in animals expressing UAS-*mre11* RNAi using the pan-hindgut driver *byn*>GAL4 (Iwaki and Lengyel, 2002). After *hs*-*I-Cre*I induction, RPA3+ foci are significantly decreased in papillar cells expressing *mre11* RNAi (**Fig. 2 I, I’, J**). These data suggest that RPA3 is recruited to papillar chromosomes in an Mre11-depedendent manner in response to DSBs.

Using other DNA break strategies, we confirmed that RPA3 recruitment and persistence in papillar cells is not specific to *hs*-*I-Cre*I-induced breaks in the repetitive rDNA. Using X-Ray irradiation (IR, 20 Gy) to induce randomly located DSBs, we also find persistent Mre11 and RPA3+ foci on papillar cell chromosomes compared to larval brain progenitors (**Fig. S1C, F; Fig. S2A, B**). *I-Cre*I targets a repetitive heterochromatic region of the chromosome and that DSB responses can vary based on chromatin state (Chiolo et al., 2011; Dialynas et al., 2019; Rong et al., 2002). To specifically address this, we tested whether a single break in a defined euchromatic locus also recruits persistent RPA3 in papillar cells. We used CRISPR-Cas9 to generate DSBs at the *rosy (ry)* locus (Carvajal-Garcia et al., 2021) in the hindgut during the endocycle. Using this method, we find that RPA3+ foci persist in papillar cells (**Fig. S2C-D**), similar to our results with *hs*-*I-Cre*I. Our results are consistent with immediate recruitment of Mre11, which recruits RPA3, after DSBs in papillar cells. Unlike checkpoint-intact neural progenitors, these repair markers persist on damaged DNA during a long period of interphase and continue into the period when these cells enter mitosis.

### Mre11 is required for acentric DNA segregation and cell survival following DNA damage

To determine the function of persistent Mre11 following DSBs in papillar cells, we induced *hs*-*I-Cre*I during the L2 stage, when papillar cells are endocycling and suppressing canonical DNA damage checkpoints **(Fig. 3A)**. Then, using *byn>GAL4* we expressed *UAS-mre11* RNAi specifically in the fly hindgut (**Fig. 3A**). Following two rounds of papillar mitoses during pupal development, four cone-shaped papillar structures made up of ∼100 cells each are present in the adult animal (Cohen et al., 2020; Fox et al., 2010). We previously found that *fancd2 RNAi* decreases adult papillar cell number specifically after DSB induction, following failure to segregate acentric DNA. In contrast, no cell number reduction occurs in *p53* null mutants or in *chk1, chk2* double mutants (Bretscher and Fox, 2016). Similar to our previous findings with *fancd2 RNAi, mre11 RNAi* adult animals contain a significantly reduced number of adult papillar cells and aberrantly shaped papillae specifically following *hs*-*I-Cre*I induction (**Fig. 3B,C**).

**Figure 3.**
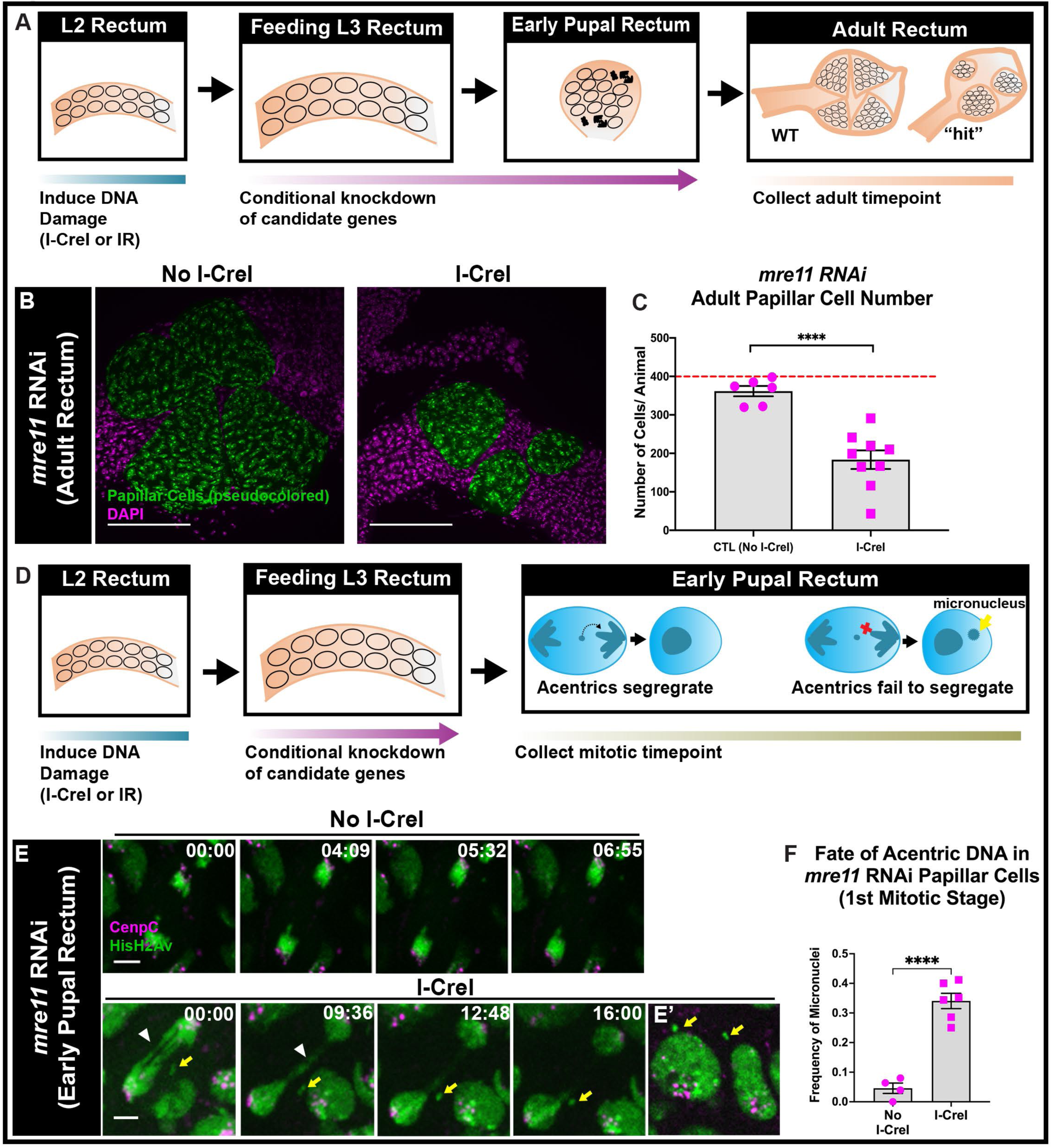
Mre11 is required for segregation of acentric DNA during mitosis. (**A**) Assay based on (Bretscher and Fox, 2016), where genes are conditionally knocked down after early development and DNA damage induction. Abnormal adult rectum phenotypes can be observed when genes important in regulating acentric DNA rescue are knocked down after DNA damage induction. (**B**) Images of adult rectum +/− *hs*-*I-Cre*I in animals expressing *mre11* RNAi in the hindgut as diagrammed in (**A**). Papillar cells (pseudo-colored), green; DNA (DAPI), magenta. Scale bars = 50μm (**C**) Quantification of adult papillar cell number in *mre11* RNAi expressing animals +/− *hs*-*I-Cre*I-induced DNA damage. Each data point represents one animal. Red dashed line indicates the expected number of papillar cells in a WT adult. Each condition has at least 2 replicates. Statistical test: Unpaired t-test, p<0.0001. (**D**) Assay based on (Bretscher and Fox, 2016), where protocol is identical to Fig 3A, except rectums are dissected during the mitotic stage where acentric fragments either properly segregate into daughter nuclei or missegregate into a micronucleus. (**E**) Live imaging of papillar cell mitosis in *mre11* RNAi animals +/− *hs*-*I-Cre*I. An acentric fragment is marked with a yellow arrow, a chromatin bridge is marked with white arrowheads. HisH2Av, green; CenpC, magenta. Scale bars = 5μm. (**F**) Quantification of the fate of acentric fragments in *mre11* RNAi papillar cells +/− *hs*-*I-Cre*I. Each condition has at least 2 replicates. Statistical test: Unpaired t-test, p<0.0001. See Methods for statistical notations.

Using fixed and *in vivo* live imaging, we followed the fate of acentric DNA during the first mitosis following DSB induction (**Fig. 3D**), using H2Av-GFP to label DNA and CenpC-Tomato to mark centromeres/kinetochores. Previously, we found that failure of acentric DNA segregation in papillar cells leads to acentric micronuclei following mitosis, as occurs in *fancd2 RNAi* animals (Bretscher and Fox, 2016). Similarly, *mre11* RNAi results in an inability to properly segregate acentric DNA during papillar cell mitosis, leading to a significant increase in acentric micronuclei (**Fig. 3E-F**). We note that, in addition to accumulating acentric micronuclei, *mre11* RNAi also results in an increase in anaphase bridges between centromere-containing chromosomes in both *hs*-*I-Cre*I (**Fig. 3E**, arrowheads) and no *hs*-*I-Cre*I animals (data not shown). This is likely due to the function of Mre11 in telomere maintenance (Syed and Tainer, 2018). To focus on acentric DNA, we only counted micronuclei that clearly lacked a Cenp-C signal, as acentric fragments lack centromeres and do not express this marker. We conclude that Mre11 is required to properly segregate acentric DNA during mitosis and for subsequent cell survival and normal papillar organ development.

### Mre11 and RPA3 have distinct localization patterns at DSBs during mitosis

As acentric fragments fail to properly incorporate during mitosis after mre*11* knockdown, we next examined Mre11 dynamics specifically during the first mitosis after DSB induction. To address this, we live imaged rectums expressing *H2Av-*RFP and *mre11-GFP* during the early (24 h post puparium formation, PPF) pupal stage. Compared to papillar cells without induced *hs*-*I-Cre*I, Mre11+ foci persist on damaged papillar chromosomes at a significantly higher frequency in these cells during early prophase as the chromosomes begin to condense (**Fig. 4A [−17:00], B**). As papillar cells with DSBs progress through mitosis, after nuclear envelope breakdown, Mre11+ foci are resolved (**Fig. 4A [00:00],B**). Therefore, Mre11+ foci are apparent at DSBs following DNA damage, persist for days on broken papillar chromosomes until early mitosis, and are resolved by nuclear envelope breakdown.

**Figure 4.**
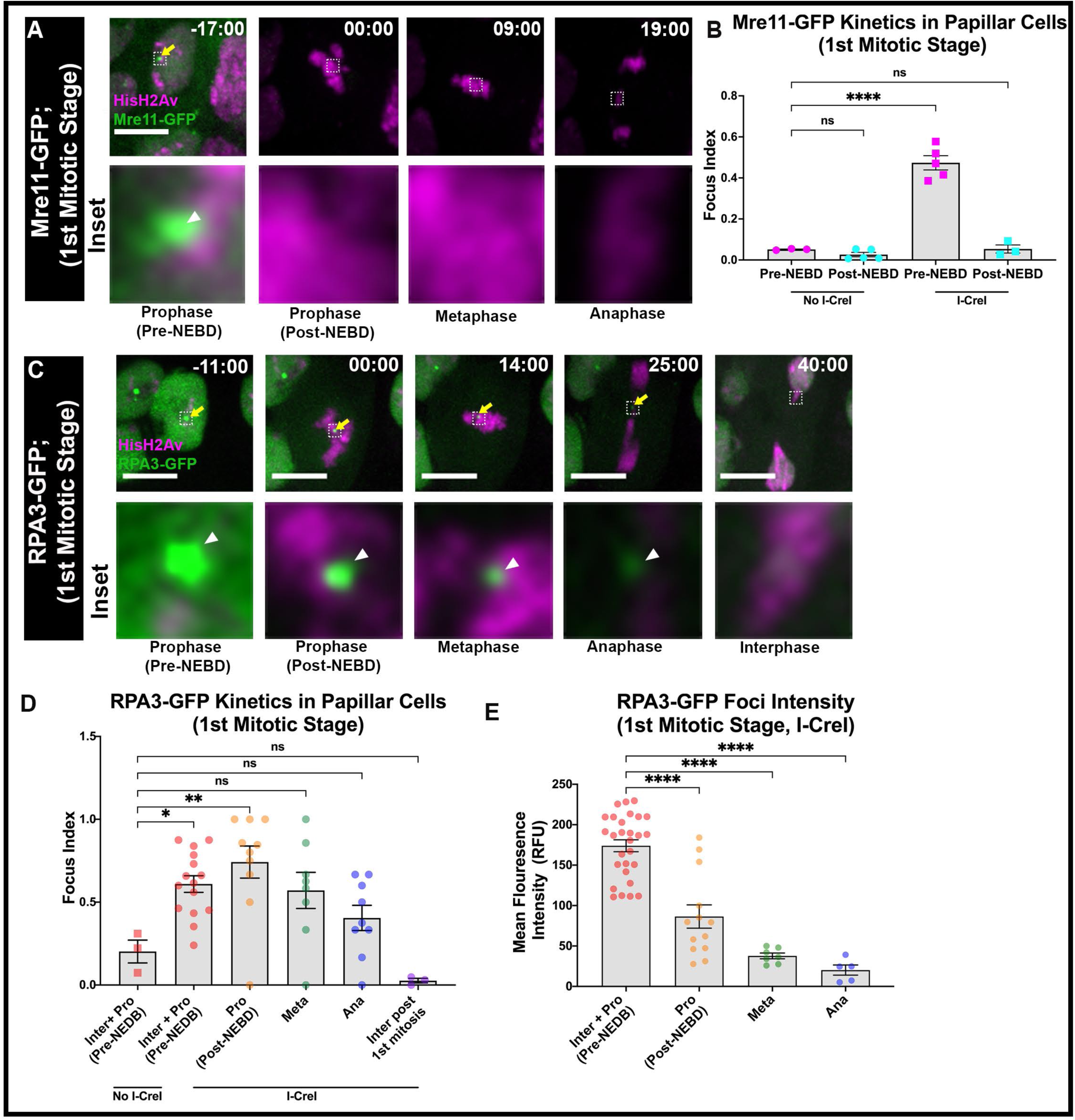
Mre11 and RPA3 have distinct kinetics during mitosis. (**A**) Mre11 localization following larval DSBs, assayed during papillar cell mitosis. Mre11+ foci are marked with yellow arrows. The hatched box highlights an area magnified 10X in the corresponding inset below each panel. Enlarged foci are marked with white arrowheads. Mre11, green; HisH2Av, magenta. Scale bars = 5μm. (**B**) Quantification of the focus index for Mre11 during the first mitotic stage in early pupal rectums. Each condition has at least 2 replicates. Statistical test: Ordinary one-way ANOVA, p<0.0001. (**C**) RPA3-GFP foci recruitment in papillar cells during mitosis +/− *hs*-*I-Cre*I. RPA3, green; HisH2Av, magenta. Labeling as in **A.** (**D**) Quantification of RPA3-GFP foci recruitment in papillar cells during mitosis +/− *hs*-*I-Cre*I. Ordinary one-way ANOVA, p= 0.0003 (**E**) Quantification of RPA3-GFP foci intensity in papillar cells during mitosis after *hs*-*I-Cre*I induction. Statistical test: Ordinary one-way ANOVA, p<0.0001. See Methods for statistical notations.

Having shown that RPA3 is recruited downstream of Mre11 (**Fig. 2I,J**), we next examined RPA3 localization kinetics during the first mitosis after induced DSBs. To address this, we live imaged papillar cells during early pupation (24 h PPF), when papillar cells are entering mitosis. Using animals expressing *H2Av-*RFP and *RPA3-*GFP, we visualized RPA3+ foci during mitosis. Similar to Mre11, RPA3+ foci are found on mitotic papillar chromosomes in a *hs*-*I-Cre*I dependent manner (**Fig. 4C [−11:00], D**). However, RPA3 differs from Mre11 in that *hs*-*I-Cre*I-dependent RPA3+ foci are found at a significantly increased level through prophase (**Fig. 4C, D**). Following prophase, the mean RPA3 focus index (the frequency in which a cell contains at least one RPA3 focus) steadily decreases (**Fig. 4D**), though a subset of cells contain RPA3+ foci that can clearly be seen on lagging DNA fragments during anaphase (**Fig. 4C, [25:00]**). RPA3+ foci are mostly resolved by the following interphase after mitosis (**Fig. 4C, [40:00]**). This gradual loss of RPA3 beginning at metaphase may reflect the onset of repair events downstream of RPA3 binding taking place between metaphase and the next interphase (see **Discussion**). We also assayed the intensity of foci, which steadily decreases as cells progress through mitosis (**Fig. 4C, E**). By the following interphase after the first papillar mitosis, the frequency of RPA3+ foci in papillar cells is significantly decreased, similar to that observed in animals without *hs*-*I-Cre*I induction (**Fig. 4C [40:00], D**). We conclude that Mre11 and RPA3+ foci persist into mitosis. While DSB-dependent Mre11+ foci are resolved prior to nuclear envelope breakdown, RPA3+ foci continue to persist at high levels until metaphase, at which point they gradually decrease on broken papillar DNA as mitosis progresses and are fully resolved by the following interphase.

### A candidate genetic screen identifies polQ as being required for acentric DNA segregation

The requirement for Mre11 and the dynamic mitotic localization of both Mre11 and RPA3 suggested that these proteins might actively signal repair events that accomplish acentric segregation. Based on our observations that Mre11 is required for RPA3 recruitment – likely due to DNA resection – we undertook a candidate genetic screen, with the hypothesis that other repair events occur downstream of RPA3 recruitment.

Using adult papillar cell number following *hs*-*I-Cre*I induction as an accessible readout to identify regulators of acentric DNA segregation during mitosis (**Fig. 3A**, Bretscher and Fox, 2016), we screened 80 candidate genes that we hypothesized to be possibly important for acentric DNA segregation (**Table S1**). We chose candidate genes based either on their known function in DNA repair or their identification in a previous cell culture-based *Drosophila* screen for DSB-responsive genes (**Table S1**, Kondo and Perrimon, 2011).

The majority of genes that we interrogated did not disrupt papillar organogenesis following DSBs. Interestingly, we did not identify a requirement for the canonical nonhomologous end-joining (cNHEJ) nor the homologous recombination (HR) repair pathways (Ceccaldi et al., 2016; Sekelsky, 2017; Wright et al., 2018). Specifically, knockdown of known regulators of cNHEJ, *ku70, ku80,* and *lig4* (Ceccaldi et al., 2015; Sekelsky, 2017), did not consistently result in a DNA damage specific decrease in papillar cell number (**Table S1**, 5/6 lines tested were not a hit). Furthermore, the requirement and recruitment of resection factors is inconsistent with cNHEJ repair, as this pathway does not require resection of DSB ends (Chang et al., 2017). Similarly, we find that Rad51, which is required for HR (Ceccaldi et al., 2015; Sekelsky, 2017; Wright et al., 2018), is not required for cell survival specifically for DNA damaged induced during the endocycle (**Table S1; Fig. S3A-B**). We do note that there is a significant decrease in cell number following *rad51* knockdown in papillar cells in the absence of *hs*-*I-Cre*I, suggesting that Rad51 plays a role in papillar organ development. However, this role is independent of *hs*-*I-Cre*I-induced DSBs, as *hs*-*I-Cre*I induction does not significantly alter adult papillar cell number in *rad51 RNAi* animals (**Fig. S3A-B**). Additionally, using an antibody for *Drosophila* Rad51, we find that Rad51 is not recruited prior to mitosis following DSB induction during the endocycle (data not shown). Rad51 foci recruitment is observed following the first mitosis, independent of exogenous DNA damage, and depends on Rad51 function (**Fig. S3C-D**), validating that we efficiently knocked down Rad51.

Our screen also included regulators of other DSB repair pathways. In contrast to the lack of requirement for c-NHEJ and HR, we identified the gene *DNA polymerase Q (polQ)* as a potential candidate for acentric DNA segregation in papillar cells. *polQ* encodes the Pol Theta protein, which is critical in the alternative end-joining repair pathway (alt-EJ) which has also been described as microhomology-mediated end-joining (MMEJ) or Theta-mediated end-joining (TMEJ) (Beagan et al., 2017; Beagan and McVey, 2016b; Chan et al., 2010; Hanscom and McVey, 2020; Kent et al., 2015; McVey and Lee, 2008). Pol Theta is involved in recognizing microhomologies near the site of a DSB and annealing homologous sequences together on either side of the break (Beagan et al., 2017; Beagan and McVey, 2016b; Chan et al., 2010; Hanscom and McVey, 2020; McVey and Lee, 2008; Sharma et al., 2015). Further, we identified several other genes involved in the alt-EJ pathway (**Table S1,** *ercc1, fen1, xpf*) as required for papillar cell survival following DSBs. As Pol Theta is a critical protein involved in several steps of alt-EJ (Beagan et al., 2017; Beagan and McVey, 2016b; Carvajal-Garcia et al., 2021; Chan et al., 2010; Liang et al., 2005; McVey and Lee, 2008; Sekelsky, 2017; Wood and Doublié, 2016)., our further analysis focused on *polQ*.

Knockdown of *polQ* using two independent RNAi lines results in both a *hs*-*I-Cre*I and IR-induced DSB-specific decrease in adult papillar cell number, leading to smaller adult papillae (**Fig. 5A,B; Fig. S3E-H**). To determine if the decrease in cell number coincides with an increase in acentric micronuclei, we observed mitotic stage papillar cells in pupae expressing UAS-*polQ* RNAi. We find a significant increase in acentric (CenpC-negative) micronuclei after *hs*-*I-Cre*I-induced DSBs in animals expressing UAS-*polQ* RNAi (**Fig 5C,D**). Then, to determine the extent to which *polQ* knockdown affects RPA3+ foci recruitment, we visualized animals expressing *ubi-RPA3-GFP* in the presence of UAS-*polQ* RNAi. We find that RPA3+ foci are present both during and prior to mitosis on broken papillar chromosomes in animals with *polQ* knockdown. However, whereas RPA3+ foci are resolved by the following interphase after mitosis in WT animals (**Fig. 4C [40:00], D**), RPA3+ foci persist into the following interphase in animals expressing UAS-*polQ* RNAi (**Fig. 5E,F**). These data suggest that our candidate screen successfully identified the alt-EJ regulator *polQ* as being required for DSB-dependent cell survival in papillar cells. *polQ* is responsible for proper acentric DNA segregation and micronuclei prevention during papillar mitosis, and for the removal of RPA3 that coincides with papillar anaphase and the following interphase.

**Figure 5:**
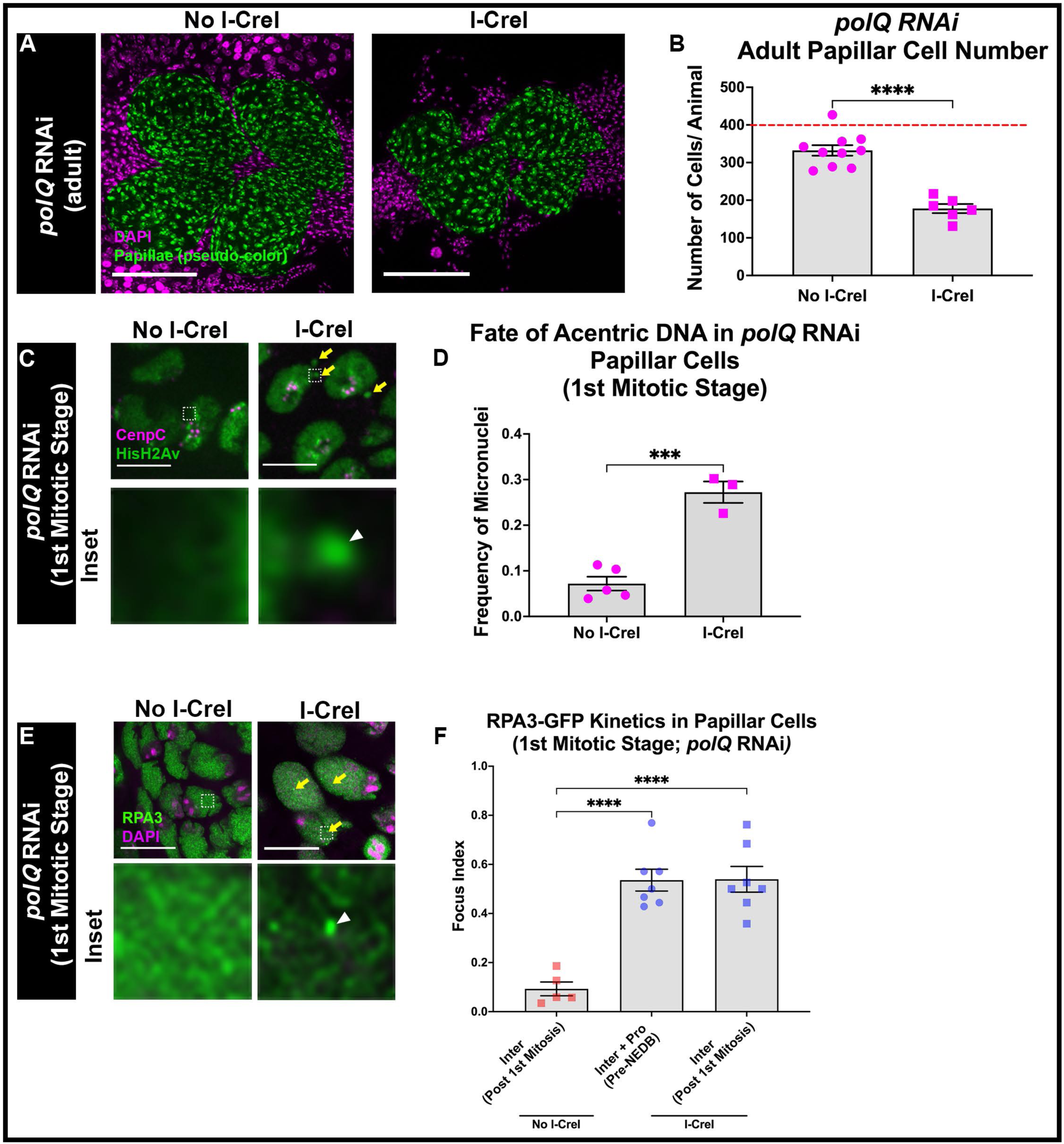
*polQ* is required for proper RPA3+ foci dynamics, acentric DNA segregation, and cell survival. (**A**) Images of adult rectum +/-*hs-I-Cre*I in animals expressing *polQ* RNAi in the hindgut. Papillar cells (pseudo-colored), green; DNA (DAPI), magenta. Scale bars = 50μm (**B**) Quantification of adult papillar cell number in *polQ* RNAi expressing animals with and without *hs-I-Cre*-induced DNA damage. Each data point represents one animal. Red dashed line indicates the expected number of papillar cells in a WT adult. Each condition has at least 2 replicates. Statistical test: Unpaired t-test, p<0.0001.(**C**) Still images of live imaged rectums expressing *polQ* RNAi +/− *hs-I-CreI*. Micronuclei are marked with yellow arrows. The hatched box highlights an area magnified 10X in the corresponding inset below each panel. Enlarged micronucleus marked with white arrowhead. HisH2Av, green; CenpC, magenta. Scale bars = 10μm. (**D**) Quantification of the fate of acentric fragments in *polQ* RNAi papillar cells +/− *hs-I-CreI*. Each condition has at least 2 replicates. Statistical test: Unpaired t-test, p=0.0003 (**E**) RPA3-GFP foci recruitment in papillar cells of *polQ* RNAi animals +/− *hs-I-CreI*. RPA3+ foci are marked with yellow arrows. The hatched box highlights an area magnified 10X in the corresponding inset below each panel. Enlarged focus marked with white arrowhead. RPA3, green; DNA (DAPI), magenta. Scale bars = 10μm. (**F**) Quantification of RPA3-GFP foci recruitment in papillar cells in *polQ* RNAi animals during mitosis +/− *hs*-*I-Cre*I. Ordinary one-way ANOVA, p<0.0001. See Methods for statistical notations.

### Monoubiquitinated Fancd2 is required for micronuclei prevention and RPA3 removal

Our analysis of *polQ* mutant papillar cells resembled our previous study of the Fanconi Anemia repair pathway protein Fancd2. This protein is also required for acentric DNA segregation during papillar cell mitosis (Bretscher and Fox, 2016). In animals expressing *UAS-fancd2* RNAi under the control of *byn*> Gal4, acentric fragments fail to properly incorporate into daughter papillar cell nuclei and instead form micronuclei (Bretscher and Fox, 2016). This increase in micronuclei is correlated with a decrease in papillar cell number and papillar organogenesis defects, which are also seen in animals lacking the Fancd2 binding partner FancI.

Fancd2 is a scaffold protein for other repair proteins that forms a heterodimer with FancI. Activation of this heterodimer and recruitment of downstream repair proteins frequently depends on monoubiquitination on conserved lysine residues in Fancd2 and FancI by the FA core complex (FancM and FancL proteins in *Drosophila*, Marek and Bale, 2006; Rodríguez and D’Andrea, 2017). To determine to what extent Fancd2 depends on its conserved monoubiquitination, we examined adult papillar cell number in FA core complex RNAi animals (**Fig. S4A-D**). Knockdown of both FA core complex members, *fancm* and *fancl*, in the hindgut using multiple independent RNAi lines, results in a *hs*-*I-Cre*I-specific decrease in papillar cell number and associated developmental defects in the adult rectum (**Fig. S4A-D**). We note that in our previous study (Bretscher and Fox, 2016) we used a *fancm* partial deletion allele (*fancm^del^*) in combination with a deficiency in the *fancm* region and observed an IR-dependent adult papillar cell phenotype but not a significant *hs*-*I-Cre*I-dependent phenotype, whereas here we find significant *hs*-*I-Cre*I-dependent papillar cell phenotypes for *fancm* with two separate RNAi lines. Taken together with our previous findings for FancI and Fancd2 (Bretscher and Fox, 2016), we conclude that the entire FA pathway is required in *Drosophila* papillar cells with DSBs.

To further test whether monoubiquitination of Fancd2 is required for papillar cell survival following DSBs, we next used two-step CRISPR-mediated homology-directed repair (Yang et al., 2020) to generate a fly where the conserved, endogenous monoubiquitination site for Fancd2, K595, is mutated to an arginine (*fancd2^K595R^,* **Fig. 6A**). This mutation has been previously observed to disrupt monoubiquitination of *Drosophila* Fancd2 *in vitro* (Marek and Bale, 2006). *fancd2^K595R^* represents the first animal model of a mutation in the endogenous conserved monoubiquitination site, and the first endogenous *fancd2* mutation of any kind in flies. *fancd2^K595R^* homozygous animals are viable and fertile, enabling us to test a role for Fancd2 monoubiquitination in acentric DNA responses. We find that in *fancd2^K595R^* animals, there is a decrease in adult papillar cell number following both *hs*-*I-Cre*I (**Fig. 6B, C**) and IR (**Fig. S4G-H**).

**Figure 6.**
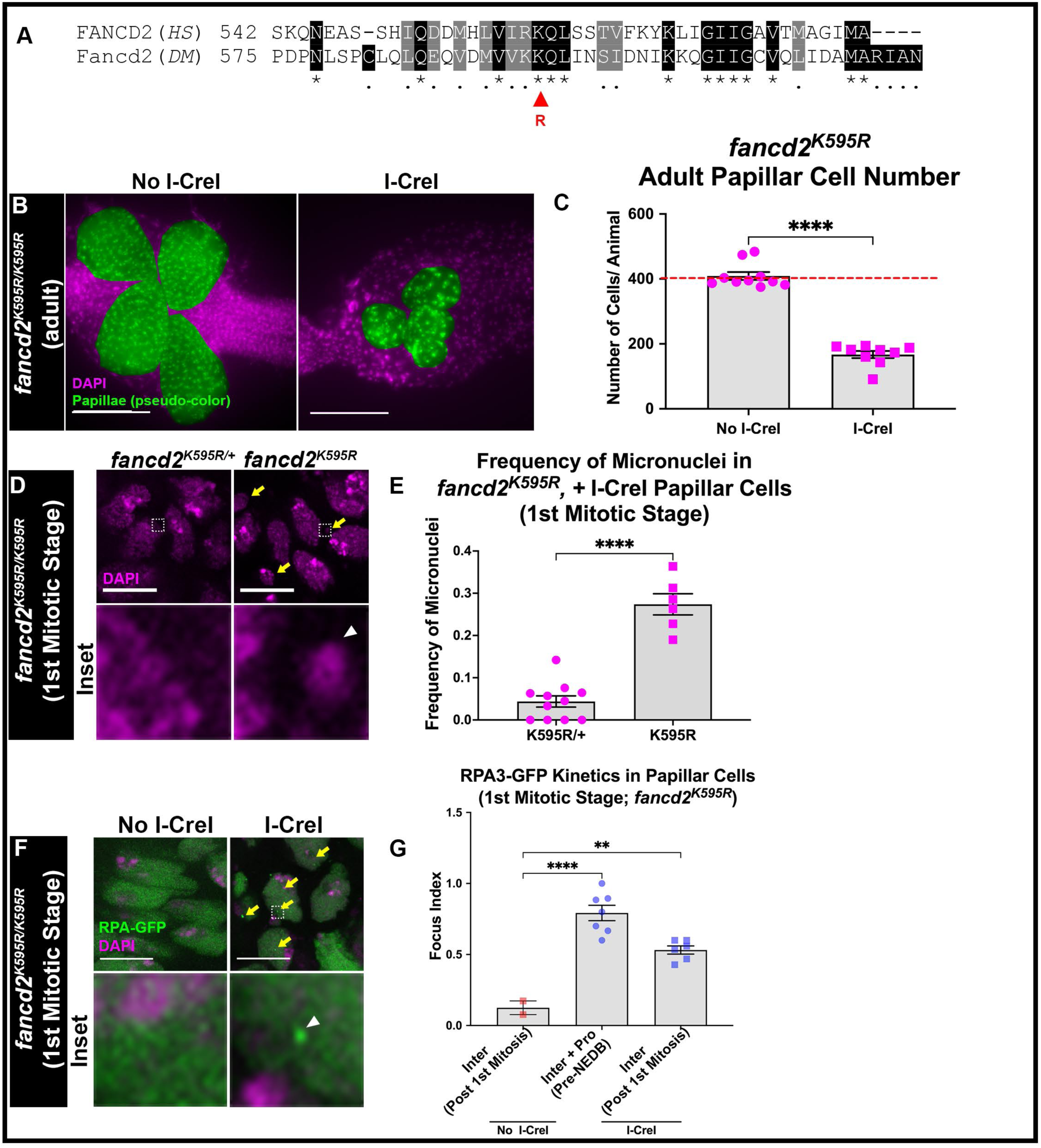
Monoubiquitinated Fancd2 is required for proper RPA3+ foci dynamics, micronuclei prevention, and cell survival. (**A**) Sequence alignment comparing the region of human (*HS*) FANCD2 protein sequence to *Drosophila* (DM) Fancd2 where the conserved monoubiquitination site is found. Asterisks and black shading indicate identical sequences whereas dots and gray shading indicate sequence similarities. Red arrowhead indicates the conserved monoubiquitinated lysine (K523 in humans; K595 in *Drosophila*). This site was mutated into an arginine (*fancd2^K595R^*) using two-step CRISPR-mediated homology-directed repair (see Methods). (**B**) Adult rectums of *fancd2^K595R^* animals +/− *hs-I-CreI*. Papillar cells (pseudo-colored), green; DNA (DAPI), magenta. (DNA). Scale bars = 50μm (**C**) Quantification of adult papillar cell number in *fancd2^K595R^* animals +/− *hs-I-CreI*. Each data point represents one animal. Red dashed line indicates the expected number of papillar cells in a WT adult. Each condition has at least 2 replicates. Statistical test: Unpaired t-test, p<0.0001. (**D**) Mitotic stage papillar cells of *fancd2^K595R^* animals +/− *hs-I-CreI*. Micronuclei are marked with yellow arrows. The hatched box highlights an area magnified 10X in the corresponding inset below each panel. Enlarged micronucleus marked with white arrowhead. DNA (DAPI), magenta. Scale bars = 10μm. (**E**) Quantification of the fate of acentric fragments with and without *hs-I-CreI-*induced DNA damage in *fancd2^K595R^* papillar cells. Each condition has at least 2 replicates. Statistical test: Unpaired t-test, p<0.0001 (**F**) RPA3-GFP foci recruitment in *fancd2^K595R^* animals +/− *hs-I-CreI*. RPA3+ foci are marked with yellow arrows. The hatched box highlights an area magnified 10X in the corresponding inset below each panel. Enlarged focus marked with white arrowhead. RPA3, green; DNA (DAPI), magenta. Scale bars = 10μm. (**F**) Quantification of RPA3-GFP foci recruitment in papillar cells in *fancd2^K595R^* animals during mitosis +/− *hs*-*I-Cre*I. Ordinary one-way ANOVA, p<0.0001. See Methods for statistical notations.

We then determined if Fancd2 monoubiquitination is required for micronuclei prevention during mitosis. To test this, we measured the frequency of micronuclei in response *hs*-*I-Cre*I DSBs. In *fancd2^K595R^* animals, the frequency of mitotic stage micronuclei is significantly higher in cells with *hs*-*I-Cre*I-induced DNA damage compared to *fancd2^K595R/+^* animals (**Fig. 6D,E**). Previous findings and the observations in this current study support a model where the FA core complex and monoubiquitination of the Fancd2-FancI heterodimer are required for micronuclei prevention during mitosis and subsequent cell survival.

We then examined whether Fancd2 monoubiquitination impacts the dynamic localization of RPA3 following DNA damage. To test this, we first visualized RPA3-GFP recruitment in animals expressing *UAS-fancd2* RNAi under the control of *byn*> Gal4. *fancd2* RNAi papillar cells fail to properly remove RPA3+ foci after mitosis following DSBs (**Fig. S4E-F**). We then visualized RPA3-GFP recruitment in *fancd2^K595R^* animals. We find that RPA3+ foci are properly recruited to papillar cells prior to mitosis in response to *hs*-*I-Cre*I-induced DSBs in *fancd2^K595R^* animals (**Fig. 6G**) and animals heterozygous for the K595R mutation (*fancd2^K595R/+^* **Fig. S4I**). However, *hs-I-Cre*I-induced DSBs increased the frequency of RPA3+ foci in *fancd2^K595R^* animals following the first mitosis (**Fig. 6F,G**). This increased frequency following mitosis is not observed in either WT (**Fig. 4C,D**) or in *fancd2^K595R/+^* animals (**Fig. S4I**). These findings suggest that, similar to Pol Theta, conserved Fancd2 monoubiquitination is required for the displacement of RPA3+ foci during mitosis.

### *polQ* and *fancd2* are epistatic for preventing micronuclei during papillar cell mitosis

Based on the observation that *polQ* and monoubiquitinated Fancd2 are both required for acentric DNA to be properly incorporated into daughter nuclei during mitosis, we next examined the genetic interaction between these two genes. FA genes have been previously implicated cooperate with genes that function in alt-EJ repair (Ceccaldi et al., 2015; Kais et al., 2016; Ward et al., 2017). Further, monoubiquitinated Fancd2 is required for recruitment of Pol Theta foci and alt-EJ in human cancer cells deficient for HR (Kais et al., 2016), which could suggest the interaction of these proteins are more important in cells lacking canonical DSB responses.

To test at a genetic level whether *polQ* and monoubiquitinated Fancd2 are working in concert to promote papillar cell survival following DNA damage, we knocked down *polQ* using UAS-*polQ* RNAi in *fancd2^K595R^* animals. After inducing *hs*-*I-Cre*I-mediated DSBs during the endocycle, we observe a significant DSB-specific decrease in adult papillar cell number in *fancd2^K595R^; polQ* RNAi animals (**Fig. 7A,B**). However, we do not observe a significant difference in adult papillar cell number after DSBs in the double mutant animals compared to either single mutant. These data suggest that *polQ* and monoubiquitinated Fancd2 work in concert to promote cell survival following DNA damage in papillar cells.

**Figure 7:**
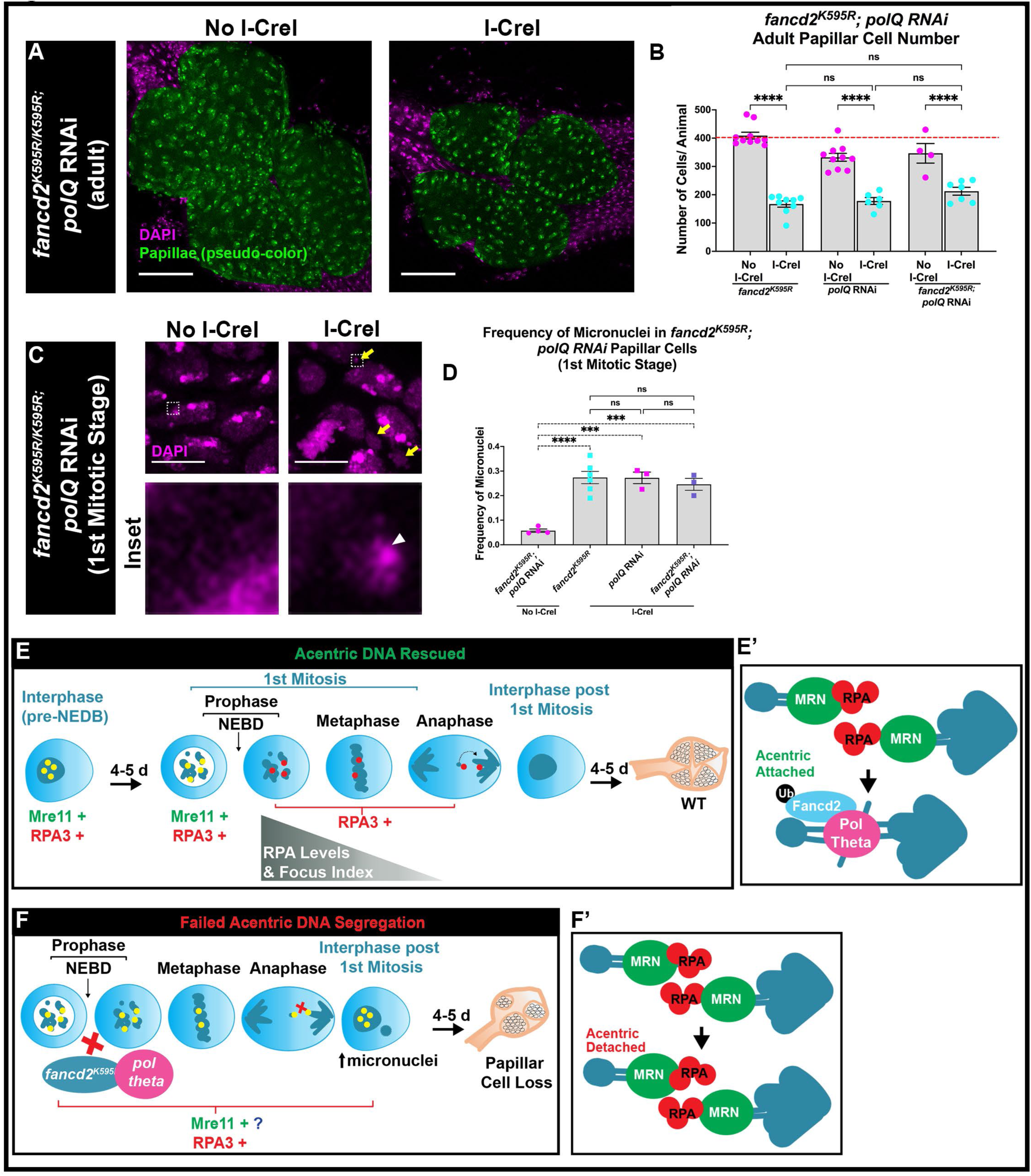
*polQ* and monoubiquitinated Fancd2 are epistatic for micronuclei prevention during papillar cell mitosis. (**A**) Images of adult rectum +/− *hs*-*I-Cre*I in animals expressing *fancd2^K595R^;polQ* RNAi in the hindgut. Papillar cells (pseudo-colored), green; DNA (DAPI), magenta. Scale bars = 50μm (**B**) Quantification of adult papillar cell number in *polQ* RNAi (data set from Fig. 5B), *fancd2^K595R^* (data set from Fig. 6C)*, fancd2^K595R^;polQ* RNAi expressing animals with and without *hs-I-CreI-*induced DNA damage. Each data point represents one animal. Red dashed line indicates the expected number of papillar cells in a WT adult. Each condition has at least 2 replicates. Statistical test: Unpaired t-test, p<0.0001 (**C**) Papillar cells at the mitotic stage in animals expressing *fancd2^K595R^;polQ* RNAi +/− *hs-I-CreI*. Micronuclei are marked with yellow arrows. The hatched box highlights an area magnified 10X in the corresponding inset below each panel. Enlarged micronucleus marked with white arrowhead. DNA (DAPI), magenta. Scale bars = 10μm. (**D**) Quantification of the fate of acentric fragments in *fancd2^K595R^;polQ* RNAi (+/− *hs*-*I-Cre*I) and *polQ* RNAi (+*hs*-*I-Cre*I) papillar cells. Each condition has at least 2 replicates. Statistical test: Ordinary one-way ANOVA, p<0.0001. (**E-F**) Model figure. (**E**) In checkpoint-inactive cells, Mre11+ and RPA3+ foci are recruited to DSBs and persist into mitosis. Mre11+ foci are resolved prior to NEBD while RPA3+ foci persist after NEDB and can be observed on lagging broken chromosomes. **(E’)** We propose a model that the requirement of monoubiquitinated Fancd2 and Pol Theta promote partial DSB repair to allow for segregation of acentric DNA during mitosis by connecting acentric DNA to centric DNA via an alt-EJ repair intermediate. (**F**) In *fancd2^K595R^* mutants and in *polQ* knockdown animals, DSB repair protein kinetics are perturbed, acentric fragments mis-segregate into micronuclei which leads to cell loss and developmental defects in the adult rectum. (**F’**) We propose a model that in *fancd2^K595R^* mutants and in *polQ* knockdown animals, resected DNA, indicated by RPA3+ foci, persists and acentric DNA is physically detached from centric DNA. These acentric fragments then mis-segregate into a micronucleus. Yellow circles= Mre11+ and RPA3+ foci; Red circles= RPA3+ foci.

We then determined the effect of *fancd2^K595R^;polQ* RNAi animals on micronuclei formation during mitosis in papillar cells following *hs*-*I-Cre*I-induced DNA damage. We observe a significant increase in the frequency of micronuclei in *fancd2^K595R^;polQ* RNAi animals after DNA damage (**Fig. 7C,D)**. Similar to our findings for adult papillar cell number, we do not observe a significant increase in micronuclei in *fancd2^K595R^;polQ* RNAi animals relative to either single gene disruption alone (*fancd2^K595R^ and polQ* RNAi, **Fig. 7D)**. This suggests that that Pol Theta and monoubiquitinated Fancd2 work in concert to prevent micronuclei during mitosis after DNA damage.

## Discussion

The extent to which damaged chromosomes can recruit DNA repair factors in the absence of a chk1/chk2/p53 DNA damage checkpoint is incompletely understood. In this current study, we find that in checkpoint-inactive papillar cells, the early acting repair proteins Mre11 and RPA3 are still recruited to DSBs despite the lack of apoptosis or a robust cell cycle arrest (**Fig. 7E**). As a consequence, Mre11 and RPA3 persist for 4-5 days following DNA damage by DSBs (**Fig. 7E**). Mre11 and RPA3+ foci are detected on mitotic chromosomes, but the kinetics of these two factors are distinct (**Fig. 7E**). Mre11+ foci are resolved prior to nuclear envelope breakdown, while RPA3+ foci persist at least into early metaphase, and in many cases are still detectable at anaphase (**Fig. 7E**). As these cells progress through mitosis, RPA3+ foci frequency and intensity significantly decrease and most cells have cleared RPA3+ foci by the following interphase (**Fig. 7E**). These dynamics are controlled by the FA and alt-EJ proteins Fancd2 and Pol Theta, respectively (**Fig. 7F**). Papillar cells that lack either Fancd2 monoubiquitination or *polQ* do not properly clear RPA3 by the end of mitosis and foci persist into the following interphase. This persistent RPA3 is accompanied by acentric DNA failing to properly segregate into daughter nuclei, leading to a significant increase in micronuclei, subsequent papillar cell loss, and adult rectum organogenesis defects (**Fig. 7F**). Based on these findings, we propose that a microhomology-mediated end joining repair intermediate links the acentric fragment to a broken, centromere containing chromosome during anaphase, thus preventing micronuclei.

### A DNA repair intermediate promotes acentric DNA segregation

The recruitment of the MRN and RPA complex components Mre11 and RPA3, respectively, to damaged papillar chromosomes, mirrors findings in other *Drosophila* cells and in mammalian cells. In *Drosophila* neural progenitors, when DSBs are induced during mitosis using micro-irradiation, Mre11 similarly localizes to broken DNA (Landmann et al., 2020). In these *Drosophila* cells, MRN is required to recruit mitotic-associated proteins such as BubR1, and repair then proceeds as these cells exit mitosis (Landmann et al., 2020). Although our prior study (Bretscher and Fox, 2016) found that papillar cells do not recruit or require BubR1 for acentric DNA segregation, our results here highlight mechanistic similarities between this neural progenitor acentric DNA response. In mammalian cells, micronuclei and chromosomal aberrations are prevented by structures known as ultra-fine DNA bridges (UFBs), which are DNA and protein based linkages that need to be resolved during mitosis (Liu et al., 2014). Such DNA-based UFB intermediates prevent micronuclei through various mechanisms. These include replication stress, which are resolved by FA proteins and BLM helicase, (Chan et al., 2007; Naim and Rosselli, 2009), or by initiated but incomplete DNA repair (Chan et al., 2018a). Several proteins that are recruited to UFBs and are required for their resolution are also either recruited or required for acentric DNA segregation during mitosis in papillar cells. The RPA complex component, RPA2, is recruited to resolved homologous recombination UFBs (Chan et al., 2018b). Similarly, we observe that RPA3 is recruited to damaged papillar chromosomes and a subset of broken DNA fragments during mitosis (**Fig. 2E-J; 4C [40:00], D**). Further, RPA2 coated UFBs are formed from BLM-induced resolution of HR intermediates, which contain Fancd2 (Chan et al., 2007; Chan et al., 2018a; Liu et al., 2014). Similarly, we previously identified BLM helicase as being required for acentric DNA segregation in damaged papillar cells (Bretscher and Fox, 2016).

We note that we have not observed long RPA3-coated UFBs during mitosis, which we hypothesize to be because papillar cells engage in alt-EJ, which can displace RPA. Alt-EJ requires annealing of short microhomologies around the site of a break, as opposed to long range end resection and strand invasion found during HR (Beagan et al., 2017; Beagan and McVey, 2016b; Chan et al., 2010; Ciccia and Elledge, 2010; Hanscom and McVey, 2020; Harper and Elledge, 2007; Jackson and Bartek, 2009; Lord and Ashworth, 2012; McVey and Lee, 2008; Sekelsky, 2017). While unwound invaded strands produce long RPA-coated strands, we speculate that alt-EJ repair displaces RPA and promotes subsequent annealing of broken DNA ends by Pol Theta and monoubiquitinated Fancd2 (**Fig. 7E’**). This unwound, annealed DNA may not be able to be detected by using reporters for histones (HisH2Av) or DAPI, but may be detectable using reporters for unwound DNA or proteins believed to be recruited downstream of RPA3 removal. Future work beyond the scope of this manuscript can investigate in detail the structure of a possible alt-EJ intermediate that tethers acentric DNA in papillar cells. As *polQ* is both required for RPA3 removal during acentric DNA segregation in papillar cells and progression of alt-EJ repair, future studies addressing if Pol Theta is recruited to acentric/centric DNA gaps will be useful (McVey and Lee, 2008). Further, full alt-EJ requires repair synthesis during gap-filling (Beagan et al., 2017; Beagan and McVey, 2016a; McVey and Lee, 2008; Sallmyr and Tomkinson, 2018). Future studies can determine the extent to which alt-EJ occurs in papillar cells, and the role of Fancd2 in this process, given the epistatic relationship that we uncovered here between *polQ* and *fancd2*.

Although alt-EJ is often described as a “back-up” pathway when major DSB repair pathways are unavailable (Iliakis et al., 2015), there is evidence that alt-EJ may be the prominent repair pathway of choice in certain conditions (Thyme and Schier, 2016). In zebrafish, alt-EJ is the DSB repair pathway of choice during early development (Thyme and Schier, 2016). It is possible that inactivation of DSB checkpoint components leads specific cell types to activate alt-EJ as the repair pathway of choice. Importantly, there is evidence that alt-EJ can occur during mitosis and during the following G1 (Deng et al., 2019; Fernandez-Vidal et al., 2014). In *Xenopus* mitotic extracts, *polQ* is important for repair of replication-induced damage specifically during mitosis, where HR and NHEJ are inactive (Deng et al., 2019). In human cells, Pol Theta binds to chromatin fractions as cells exit mitotis and enter G1, even in the absense of DNA damage (Fernandez-Vidal et al., 2014). Taken together, these findings support a model in which Pol Theta can function during mitosis to displace RPA3 and promote alt-EJ of acentric DNA ends.

### Fancd2 monoubiquitination and Pol Theta work together to promote acentric DNA segregation

Our data support a model in which the monoubiquitination of Fancd2 and Pol Theta cooperate to promote acentric DNA segregation during papillar cell mitosis. Mechanisms for how monoubiquitinated Fancd2 and Pol Theta cooperate to promote repair has been recently characterized (Kais et al., 2016; Muzzini et al., 2008). In *C. elegans*, Pol Theta is required for FA-dependent inter-strand cross-link repair (Muzzini et al., 2008). At stalled replication forks, monoubiquitinated Fancd2 is required for Pol Theta foci formation and for efficient alt-EJ repair (Kais et al., 2016; Patel et al., 2021). Additionally, Fancd2 and FEN1 (hit in our screen, **Table S1**), an endonuclease involved in alt-EJ, are epistatic in the repair of chemotherapy (PARP inhibitor and cisplatin)-induced DNA damage (Ward et al., 2017). We propose a model where, in checkpoint-inactive papillar cells, monoubiquitination of Fancd2 by the FA core complex leads to the recruitment of Pol Theta to resected, RPA coated-ssDNA. Pol Theta is then responsible for displacing RPA and promoting alt-EJ, allowing for an alt-EJ repair intermediate to form between acentric DNA and its centric partner promoting proper segregation of acentric DNA to daughter nuclei (**Fig**. **7E’**). When Fancd2 monoubiquitination is disrupted or when *polQ* is knocked down, RPA-coated resected DNA persists and acentric DNA remains detached from centromere containing chromosomes. This then leads to mis-segregation of acentric DNA into the formation of micronuclei (**Fig. 7F**’).

### Papillar cell acentric DNA segregation as a model for cancer cell biology and tumor resistance

We have previously described characteristics of papillar cell biology that are similar to what is observed in cancer cells (Cohen et al., 2020; Peterson et al., 2020). Papillar cells undergo endocycles during the second larval instar stage (Fox and Duronio, 2013; Øvrebø and Edgar, 2018). Over one-third of cancers experience whole genome duplication events, making polyploidy one of the most common genetic alterations in cancer (Zack et al., 2013). Additionally, papillar cells are resistant to high levels of X-ray radiation (20 Gy) and other forms of DNA damage (Bretscher and Fox, 2016). Not only do cancers cells show resistance to various therapies that cause DNA damage such as chemotherapy drugs or irradiation therapy, cancer cells that have undergone genome duplication events also show an increase in resistance to therapies as well (Shen et al., 2008; Shen et al., 2013; Szakács et al., 2006). One potential mechanism for how cancer cells and papillar cells tolerate high levels of DNA damage may be due to a dysregulated DDR (Bretscher and Fox, 2016). Several cancers including in the context of whole genome duplications have some degree of DDR disfunction to promote a proliferative advantage and genome instability (Lord and Ashworth, 2012; Zheng et al., 2012). As highlighted here, our findings are relevant to prevention of micronuclei. Micronuclei are linked to a variety of different disease phenotypes and cancer (Marcozzi et al., 2018). The presence of micronuclei can lead to chromothripsis, a genome shattering event characterized by deleterious chromosome rearrangements (Crasta et al., 2012; Nazaryan-Petersen et al., 2020; Zhang et al., 2015). Micronuclei can also illicit an innate immune response through the cGAS-STING pathway (Liu et al., 2018; McLaughlin et al., 2020). This micronuclei-induced inflammatory response has been linked to tumor formation and metastasis (Bakhoum et al., 2018; Harding et al., 2017; Mackenzie et al., 2017).

In this current study we have added to the similarities between papillar cell and cancer cell biology. Pol Theta and Fancd2 are highly upregulated in HR-deficient cancers (Ceccaldi et al., 2015; D’Andrea, 2018; Kais et al., 2016; Wood and Doublié, 2016). We find that papillar cells fail to recruit HR regulator Rad51 in response to induced DSBs from the time of polyploidization (endocycle) to the first round of mitotic division (**Fig. S3C,D**). Furthermore, papillar cell survival following DSBs does not depend *on rad51* (**Fig. S3A,B**). Instead, papillar cells rely on both *polQ* and monoubiquitinated Fancd2 for proper segregation of acentric DNA during mitosis. It is therefore possible that papillar cells represent an *in vivo* model for understanding HR-deficient cancer cell biology. This will allow us to further understand the mechanisms of Fancd2 and Pol Theta in HR-deficient tumors as well as investigate novel gene targets in treatment-resistant cancers. Although there is great promise for chemotherapies that target HR-deficient tumors (e.g. PARP inhibitors), these drugs are highly cytotoxic as PARP is important for various cellular processes (Ricks et al., 2015) and cancer cells can develop PARP inhibition resistance (Ceccaldi et al., 2016; Jenkins et al., 2012; Patel et al., 2021). Understanding how to target Fancd2 monoubiquitination, improve *polQ* inhibition, and the identification of novel targets would be of great interest in addressing cancer therapy resistance.

## Materials and Methods

### *Drosophila* Stocks

A detailed list of stocks used in this study is available in **Table S2**.

### *Drosophila* Culture & Genetics

Flies were raised on standard fly food (Archon Scientific, Durham, NC) at 22°C except for when experiments were conducted with *tub-Gal80^ts^.* In experiments using *tub-Gal80^ts^*, animals were raised at 18°C until the second larval instar stage, when DNA damage was induced (see **DNA Damage**). Those animals were then shifted to 29°C during the feeding third larval instar stage.

The *fancd2^K595R/K595R^* flies were generated by GenetiVision Corporation (www.genetivision.com). The K595R mutation was created via two steps of CRISPR-Cas9 mediated HDR (homology directed repair) events. In the first step, two CRISPR-Cas9 targets were designed to delete a 490 bp fragment containing the K595. Two guide RNAs were cloned into pCFD3 vector (http://www.flyrnai.org/tools/grna_tracker/web/files/Cloning-with-pCFD3.pdf) and a donor DNA was created with a GFP cassette flanked by two 1 kb f*ancd2* sequences beyond cleavage sites. Upon co-injection of both DNA constructs, two gRNAs were expressed to direct the double strand break (DSB) by Cas9 (endogenously expressed in the injection stock BL#54591). After the DSB, the GFP cassette was inserted into *fancd2* genome via donor DNA mediated recombination. In the second step, based on the same principle, a 1039 bp DNA containing GFP cassette (plus neighboring *fancd2* sequences) was substituted by the mutant allele containing K595R using a new set of gRNAs and new donor DNA which is the *fancd2* sequence with K595R mutation introduced.

The stock containing *ry^776+^* gRNA (ATTGTGGCGGAGATCTCGA) was made by the Sekelsky lab at UNC Chapel Hill, NC. The guide RNA was cloned into a pCFD3 vector (Fillip Port, Simon Bullock lab, MRC-LMB) and injected at 58A using PhiC 31. The landing site has a 3xP3 DsRed marker and the pCFD3 has a *vermillion+* marker.

### DNA Damage

Double-strand breaks (DSBs) were induced using three methods. The first method uses animals expressing the *hs*-*I-Cre*I transgene. DNA damage was induced using methods in (Bretscher and Fox, 2016). Animals were heat shocked at 37°C for 90 min at the second larval instar stage. The second method to induce DSBs was using IR as described in (Bretscher and Fox, 2016). Fly food containing second larval instar flies were placed in 60 mm petri dishes. The petri dishes were then place in an X-RAD 160 PXI precision X-ray irradiator (calibrated by a dosimetrist) at 20 Gy. Lastly, DSBs were induced at the *ry* locus using CRISPR-Cas9 (see ***Drosophila* Culture and Genetics**). Animals expressing the *ry* gRNA were crossed to animals expressing *UAS-Cas9*; *byn-Gal4, tub-Gal80ts*. These animals were raised at 18°C until the second larval instar stage. Larvae were then shifted to 29°C to allow for expression of Cas9 and subsequent induction of DSBs at the *ry* locus.

### Fixed Imaging

*Drosophila* tissues were dissected in 1X PBS and immediately fixed in 3.7% formaldehyde + 0.3% Triton-X. Immunofluorescence (IF) staining was performed as in (Sawyer et al., 2017). Primary antibodies used in this study were mouse anti-gamma H2Av (1:2,500, Lake et al., 2013), rabbit anti-GFP (1:1,000, Life Technologies), and rabbit anti-Rad51 (A gift from J. Kadonaga, USCD). Tissue was stained with DAPI at 5 μg/ml.

All fixed images presented in figures were acquired were acquired using Zeiss 880 Airyscan Inverted Confocal using 20x/0.80 (420650-9901): Plan-Apochromat, NA: 0.80, air and 63x/1.4 (420782-9900): Plan-Apochromat, NA: 1.4, Oil objectives. 405 nm Diode, Argon/2 488, and 561 nm Diode lasers were used on a Zeiss Axio Observer Z1 with Definite Focus2. The system was controlled by Zeiss Zen 2.3. Adult rectum images were acquired using Zeiss AxioImager M.2 with Apotome processing using 20x objective. Additional imaging used for quantitation only were performed using Andor Dragonfly 505 unit with Borealis illumination spinning disk confocal with Andor iXon Life 888 1024×1024 EMCCD camera and using 63x/1.47 TIRF HC PL APO CORR (Leica 11506319) Oil objective (405 nm (100 mw), 488 nm (150 mW), and 561 nm (100 mW) diodes) on a Leica DMi8 microscope. The system was controlled by Fusion software.

### Live Imaging

Tissues were prepared for live imaging as described in previous studies (Fox et al., 2010; Schoenfelder et al., 2014) Images were acquired on spinning disk confocal (Yokogawa CSU10 scanhead) on an Olympus IX-70 inverted microscope using a 60×/1.3 NA UPlanSApo Silicon oil, 488 and 568 nm Kr-Ar laser lines for excitation, and an Andor Ixon3 897 512 electron-multiplying charge-coupled device camera. The system was controlled by MetaMorph 7.7

### Image Analysis

All image analysis was performed using ImageJ (Schneider et al., 2012). Focus index was determined as the percentage of cells with at least one DNA repair focus. Mre11-GFP foci in fixed tissues were not as distinguishable as RPA3-GFP foci and thus were determined using co-localization with gamma H2AV antibody. Focus index, micronuclei frequency, and adult papillar cell number were quantified using Image J Cell Counter plugin. Foci Intensity was calculated by measuring mean fluorescence intensity of individual foci. Images were converted to grayscale and individual ROIs were drawn around multiple foci within each cell across different stages of cell division (interphase, prophase (pre and post NEBD), metaphase, anaphase, and interphase following the first division). Measurements of individual ROIs were taken using the mean gray value. ROIs of the same size were taken from background on the same stack and timepoint as the relevant foci and mean intensity values were averaged and subtracted from each focus mean intensity value.

### Statistics

All statistics were computed in Prism (versions 8 and 9, GraphPad, La Jolla, CA). Adult papillar cell number and micronuclei frequency for animals expressing various RNAi constructs were analyzed using an unpaired t test (no DNA damage versus DNA damage). Focus index and fluorescence intensity measurements were analyzed using ordinary one-way ANOVA with multiple comparisons (data points compared to no DNA damage control). P-values and N values are indicated in figure legends. Statistical notations used in figures: NS= not significant, *=P ≤ 0.05, **= P ≤ 0.01, ***=P ≤ 0.001, ****= P ≤ 0.0001.

## Acknowledgments

The following kindly provided reagents used in this study: Bloomington Drosophila Stock Center, Developmental Studies Hybridoma Bank, Vienna Drosophila Resource Center. We thank Fu Yang, PhD and Zhao Zhang, PhD for comments on the manuscript. This project was supported by NIGMS grant GM118447 to D.T.F., an NSF GRFP grant to D.E.C., and NCI grant F31CA186545 to H.S.B. We thank Lisa Cameron and Yasheng Gao at the Duke Light Microscopy Core Facility for assistance.

## Author contributions

Conceptualization: D.E.C., H.S.B., D.T.F.; Methodology: D.E.C., H.S.B., E.A.J., K.B.B. Validation: D.E.C., H.S.B., E.A.J., K.B.B.; Formal Analysis: D.E.C., H.S.B., E.A.J., K.B.B., D.T.F.; Investigation: D.E.C., H.S.B., E.A.J., K.B.B.; Data curation: D.E.C., H.S.B., E.A.J., K.B.B.; Writing: D.E.C., D.T.F.; Visualization: D.E.C., H.S.B., E.A.J.; Supervision: D.T.F.; Funding acquisition: D.E.C., H.S.B., D.T.F.

## Supplementary Material

### Supplemental Figure Legends

**Supplemental Table 1.**
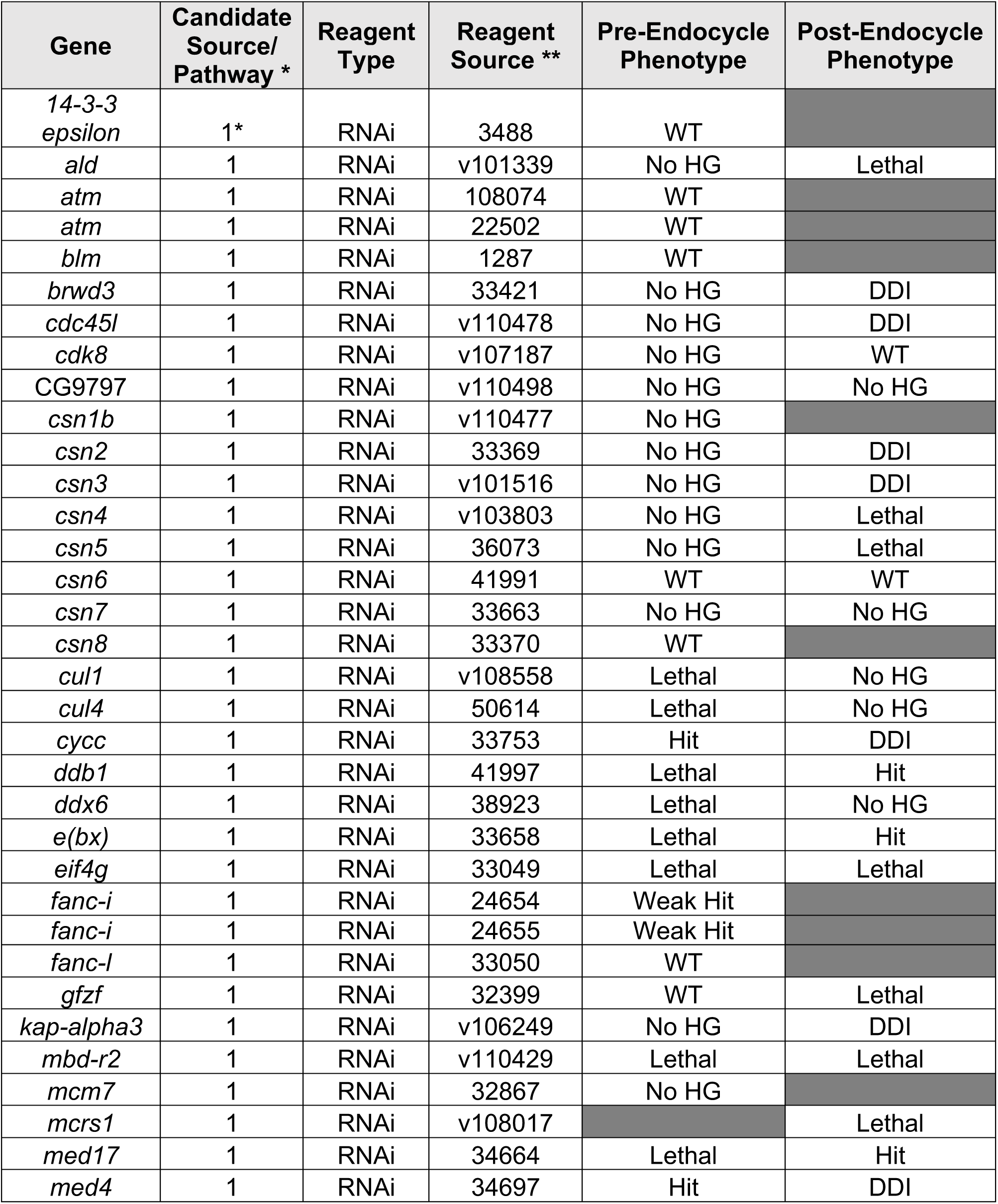
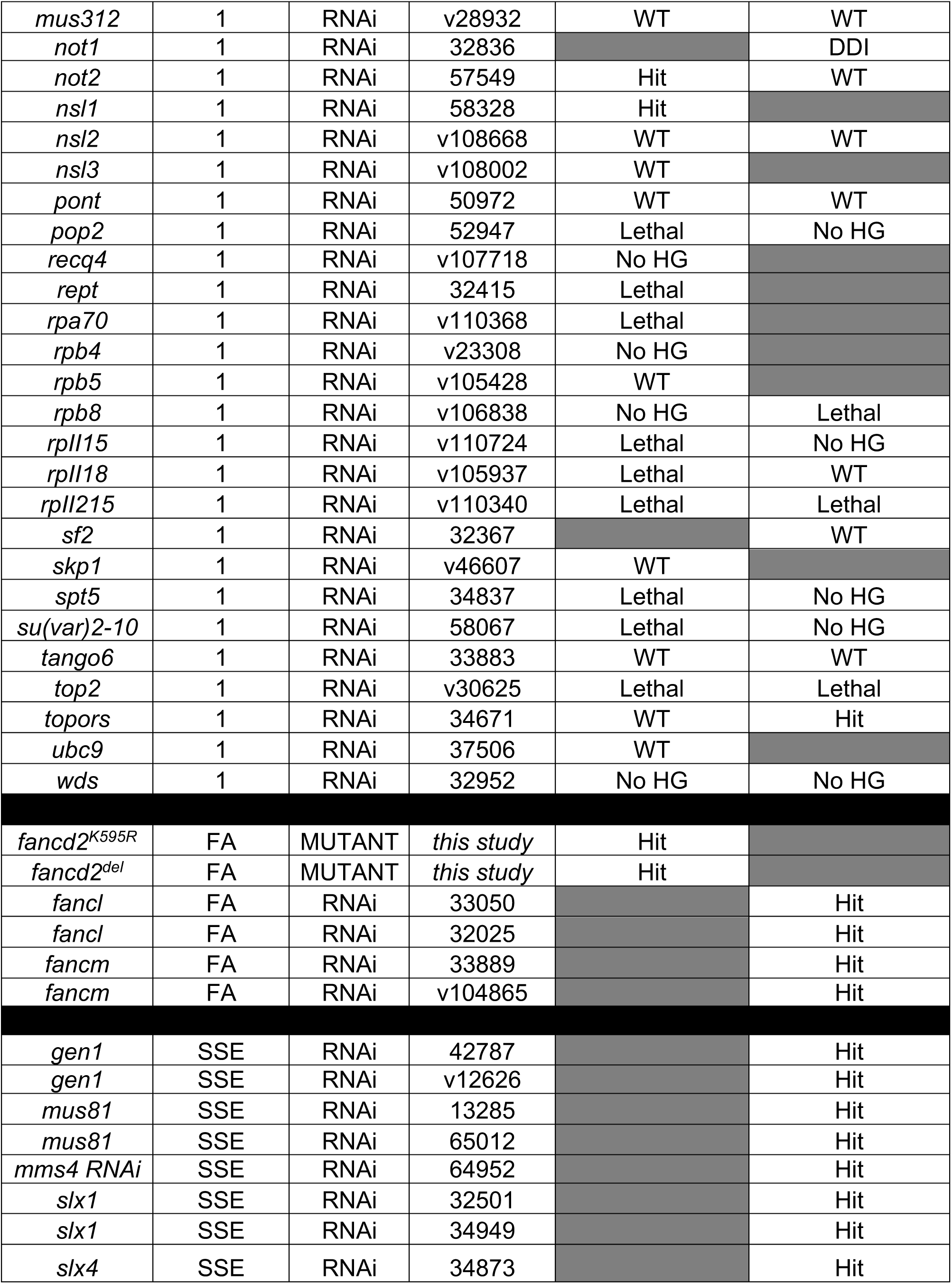
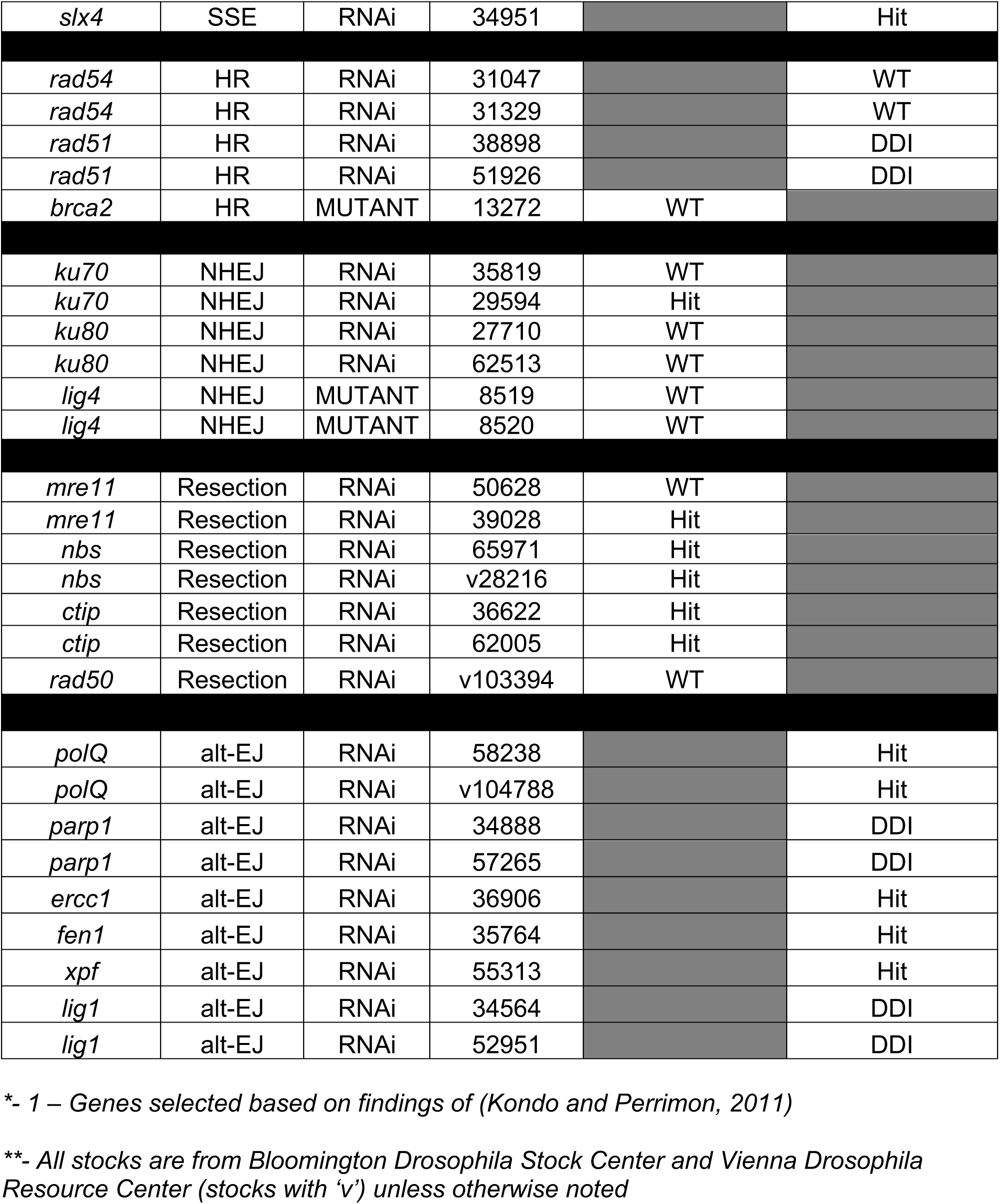
Candidate Screen for acentric DNA segregation in papillar cells. A candidate screen of 80 genes was performed to identify regulators of acentric DNA regulation in papillar cells. See **Fig. 3A** for details of screen. Wildtype (WT) - expected adult papillar cell number (∼400) is observed. Hit - average adult papillar cell number is significantly lower than expected with DNA damage. No HG - No hindgut. DDI - DNA damage-independent decrease in papillar cell number. Pre-Endocycle Phenotype - RNAi was activated prior to the endocycle using byn>Gal4 or mutant construct was used. Post-Endocycle Phenotype - RNAi was activated after the endocycle using byn>Gal4, tub-Gal80TS. SSE – Structure Specific Endonuclease

**Supplemental Table 2.**
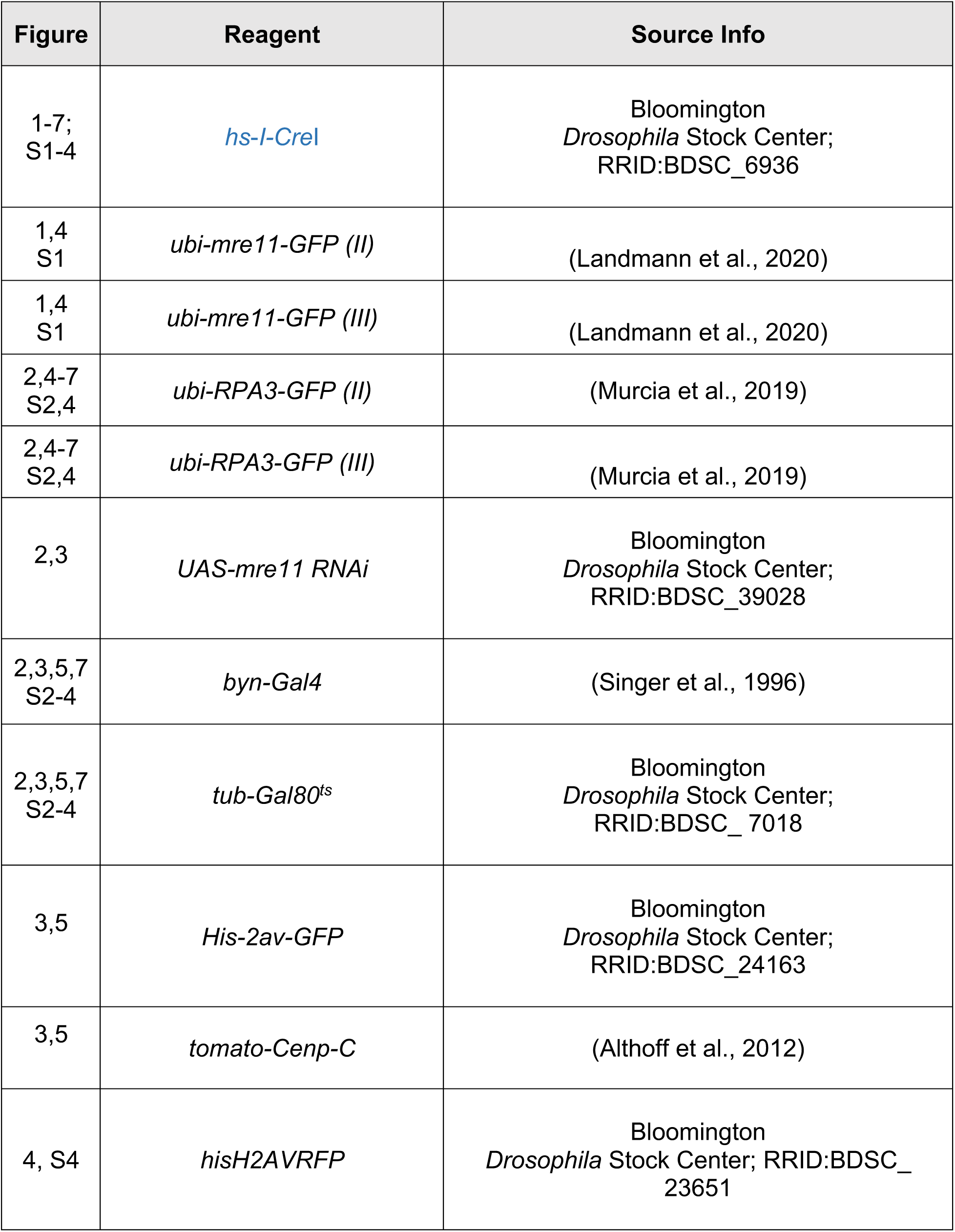
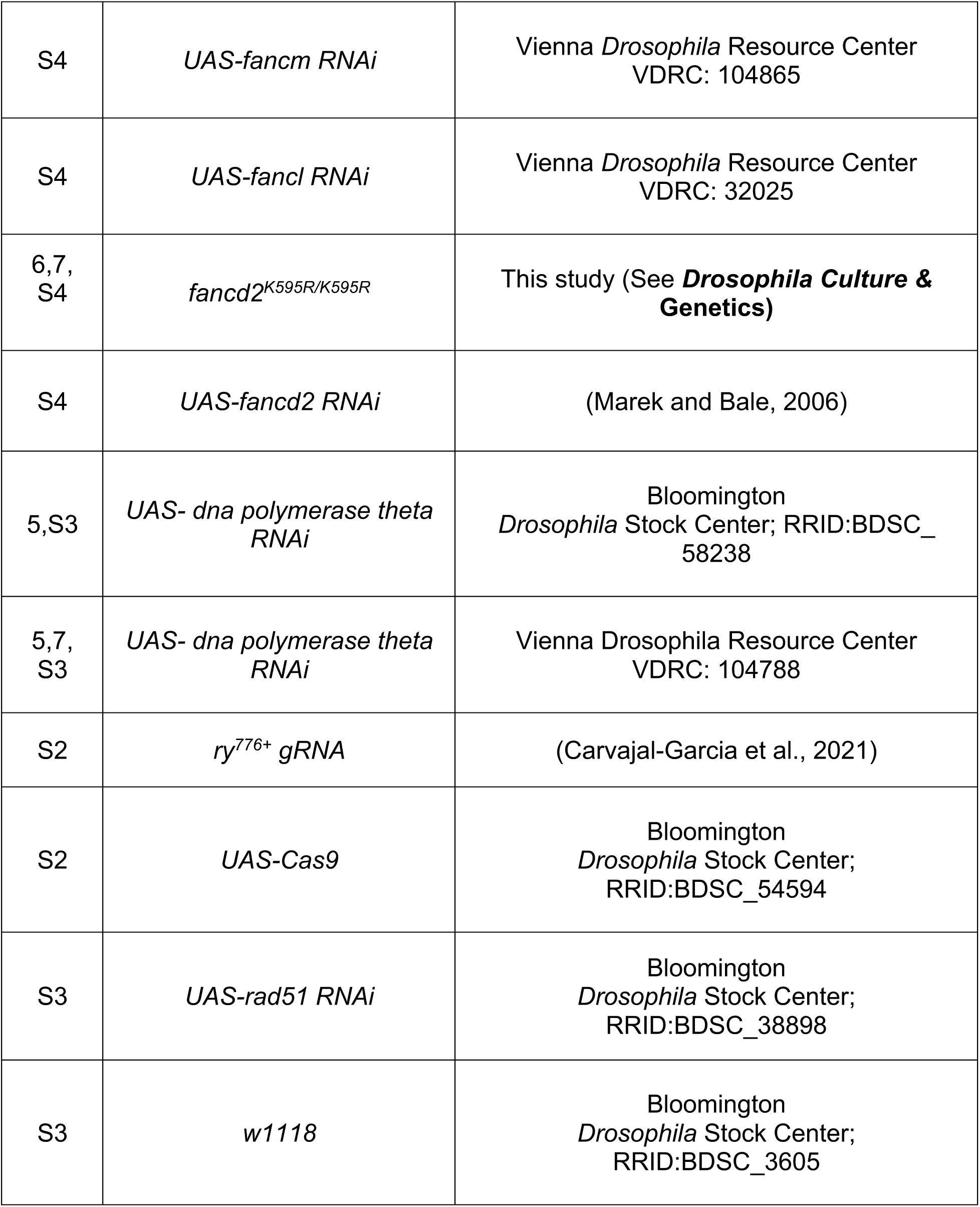
Fly stocks used in this study. This table lists all fly stocks used in this study, the source, and figures in which they were used.

**Supplemental Figure 1.**
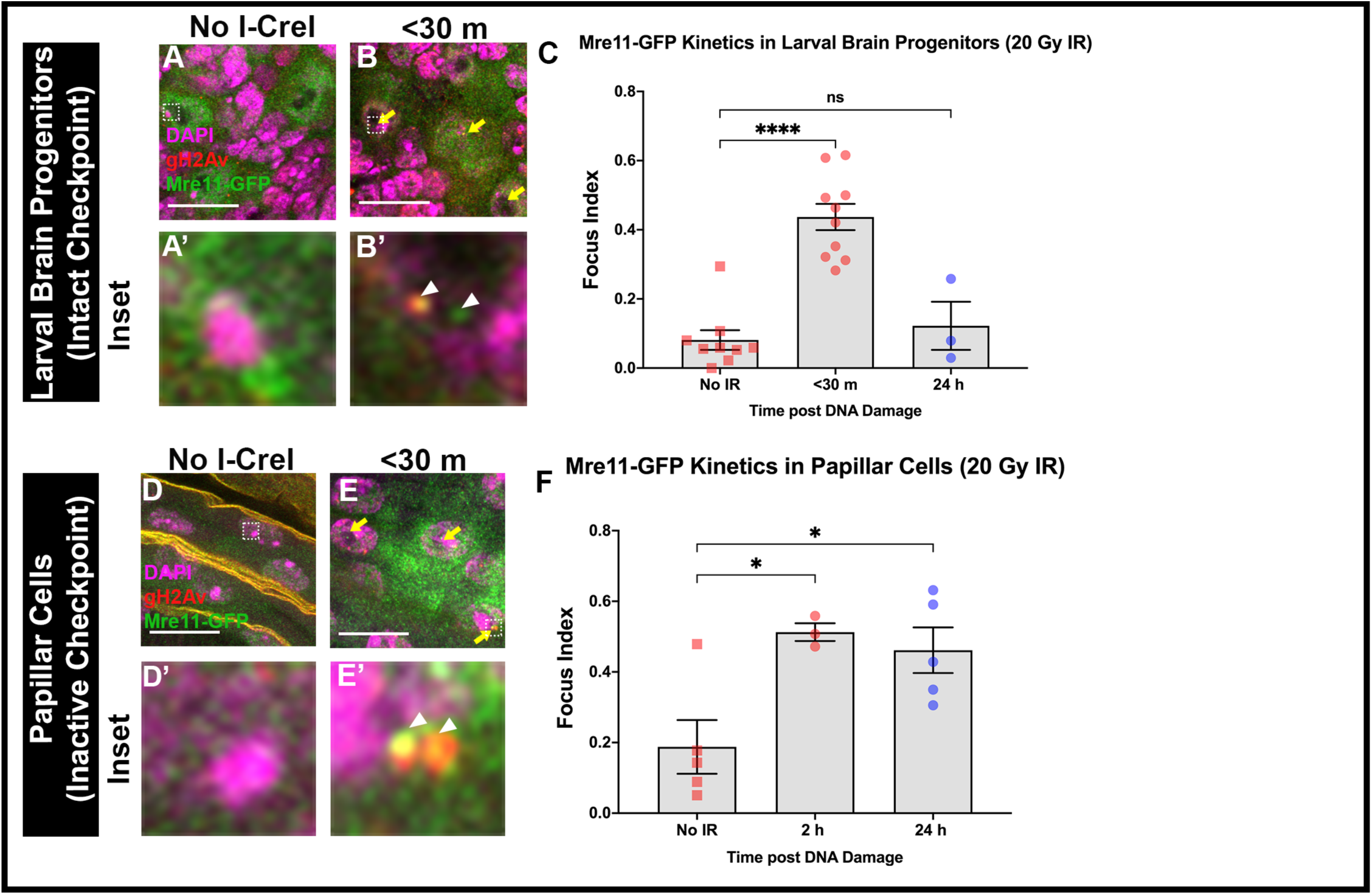
Mre11 colocalizes with *γ*H2Av and is recruited to papillar cells with delayed kinetics after IR-induced DSBs. (**A-B’**) Mre11-GFP co-localization with !H2Av in larval brain progenitors less than 30 m after +/− *hs*-*I-Cre*I. Time after break induction is indicated in minutes (m). Mre11+ foci are marked with yellow arrows. The hatched box highlights an area magnified 10X in the corresponding inset below each panel. Enlarged foci are marked with white arrowheads. (**C**) Mre11-GFP localization over time in larval brain progenitors +/− IR. Each data point represents one animal. Each timepoint has at least 2 replicates. Statistical test: Ordinary one-way ANOVA, p<0.0001. (D-E’) Mre11-GFP co-localization with !H2Av in papillar cells less than 30 m after +/− *hs*-*I-Cre*I. Labeling as in **A-B’**. (**F**) Mre11-GFP localization over time in papillar cells +/− IR. Each data point represents one animal. Each timepoint has at least 2 replicates. Statistical test: Ordinary one-way ANOVA, p=0.0147. See Methods for statistical notations.

**Supplemental Figure 2.**
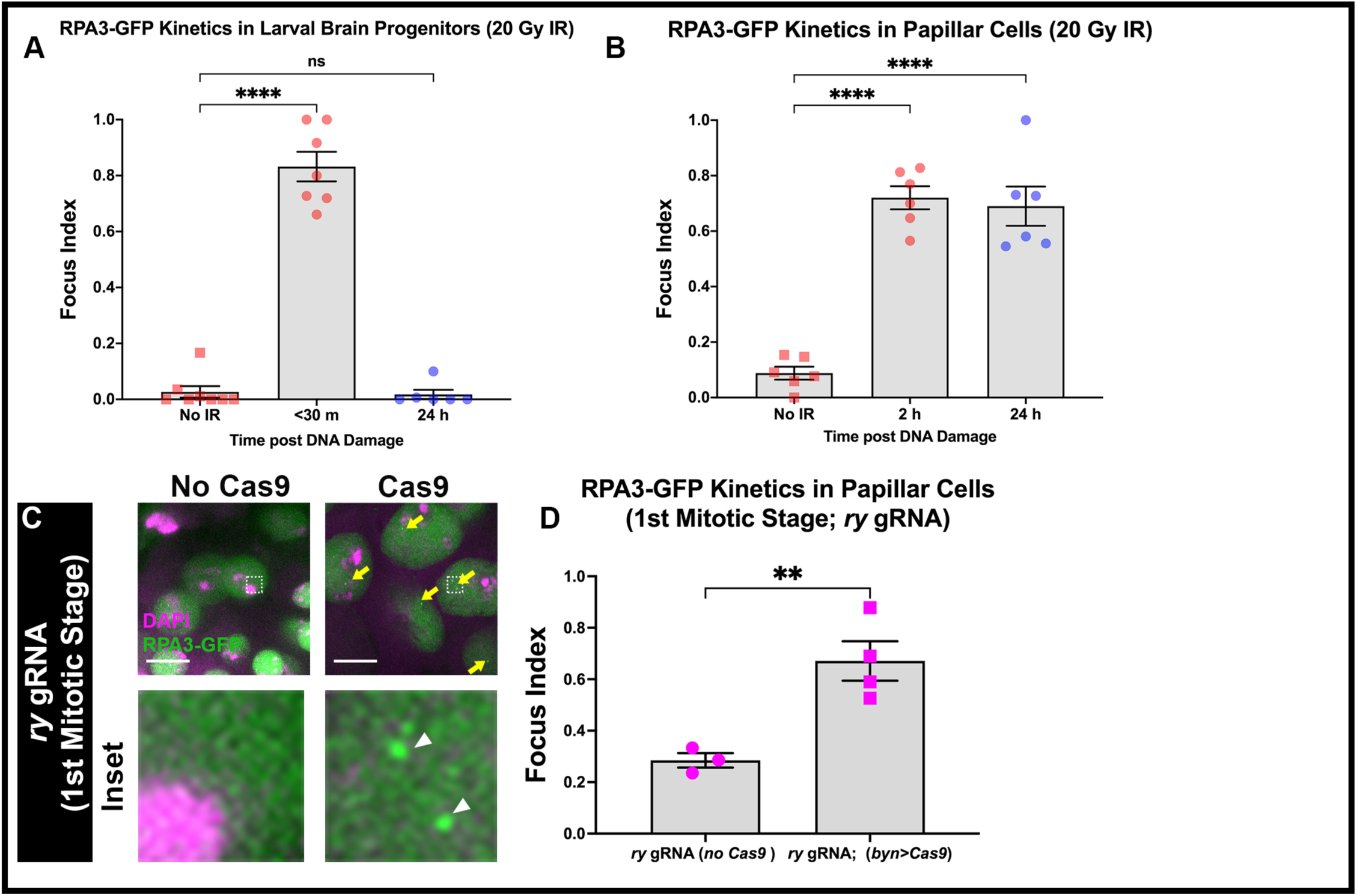
RPA3 is recruited to papillar cells with delayed kinetics after IR-induced DSBs. (**A**) RPA3-GFP localization over time in larval brain progenitors +/− IR. Each data point represents one animal. Each timepoint has at least 2 replicates. Statistical test: Ordinary one-way ANOVA, p<0.0001. (**B**) RPA3-GFP localization over time in papillar cells +/− IR. Each data point represents one animal. Each timepoint has at least 2 replicates. Statistical test: Ordinary one-way ANOVA, p<0.0001. See Methods for statistical notations. (**C-D**) RPA3-recruitment to papillar cells +/− Cas9-induced DSBs using *ry* gRNA. RPA3+ foci are marked with yellow arrows. The hatched box highlights an area magnified 10X in the corresponding inset below each panel. Enlarged foci are marked with white arrowheads. (**D**) Quantification of RPA3-GFP foci recruitment in papillar cells in papillar cells with +/− Cas9-induced DSBs using *ry* gRNA during the 1^st^ mitotic stage. Ordinary one-way ANOVA, p =0.0092. See Methods for statistical notations.

**Supplemental Figure 3.**
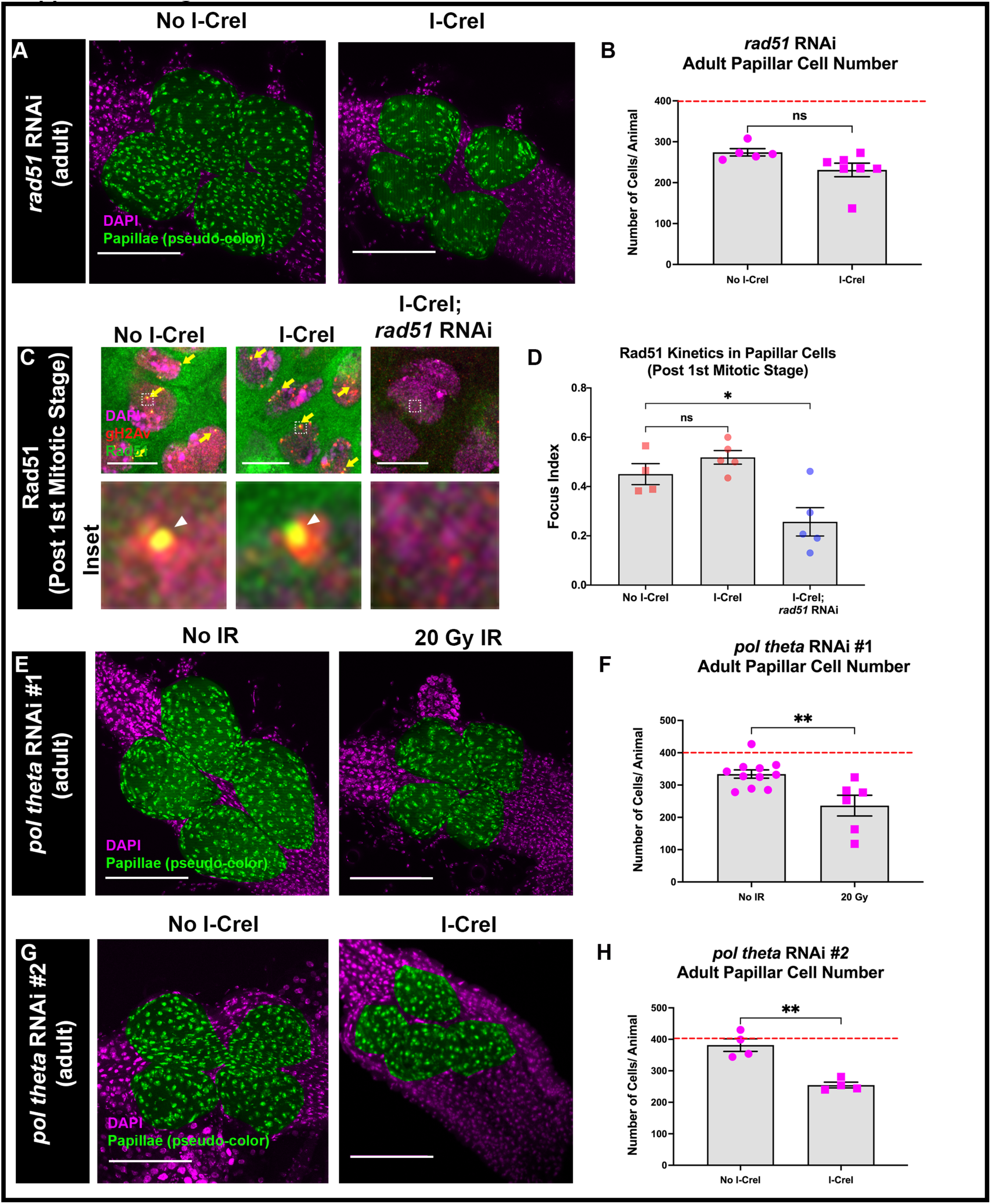
Rad51 is not required for cell survival following DSBs in papillar cells. (**A**) Adult rectums of *rad51* RNAi animals +/− *hs-I-CreI*. Green= pseudo-colored rectal papillae, magenta= DAPI (DNA). Scale bars = 100 μm (**B**) Quantification of adult papillar cell number in *rad51* RNAi animals +/− *hs-I-CreI*. Each data point represents one animal. Red dashed line indicates the expected number of papillar cells in a WT adult. Each condition has at least 2 replicates. Statistical test: Unpaired t-test, p=0.0702. (**C**) Rad51 recruitment to WT papillar cells +/− *hs*-*I-Cre*I and *rad51* RNAi papillar cells. Rad51+ foci are marked with yellow arrows. The hatched box highlights an area magnified 10X in the corresponding inset below each panel. Enlarged foci are marked with white arrowheads. (**D**) Quantification of Rad51 foci recruitment in papillar cells in +/− *hs*-*I-Cre*I and *rad51* RNAi papillar cells post the 1^st^ mitotic stage. Ordinary one-way ANOVA, p =0.0037. (**E**) Adult rectums of *polQ* RNAi animals +/− IR. Green= pseudo-colored rectal papillae, magenta= DAPI (DNA). Scale bars = 100 μm (**F**) Quantification of adult papillar cell number in *polQ* RNAi animals +/− IR. Each data point represents one animal. Red dashed line indicates the expected number of papillar cells in a WT adult. Each condition has at least 2 replicates. Statistical test: Unpaired t-test, p=0.0043 (**G**) Adult rectums of *polQ* RNAi #2 (2^nd^ RNAi line) animals +/− *hs-I-CreI*. Green= pseudo-colored rectal papillae, magenta= DAPI (DNA). Scale bars = 100 μm (**H**) Quantification of adult papillar cell number in *polQ* RNAi #2 animals +/− *hs-I-CreI*. Each data point represents one animal. Red dashed line indicates the expected number of papillar cells in a WT adult. Each condition has at least 2 replicates. Statistical test: Unpaired t-test, p=0.0012. See Methods for statistical notations.

**Supplemental Figure 4.**
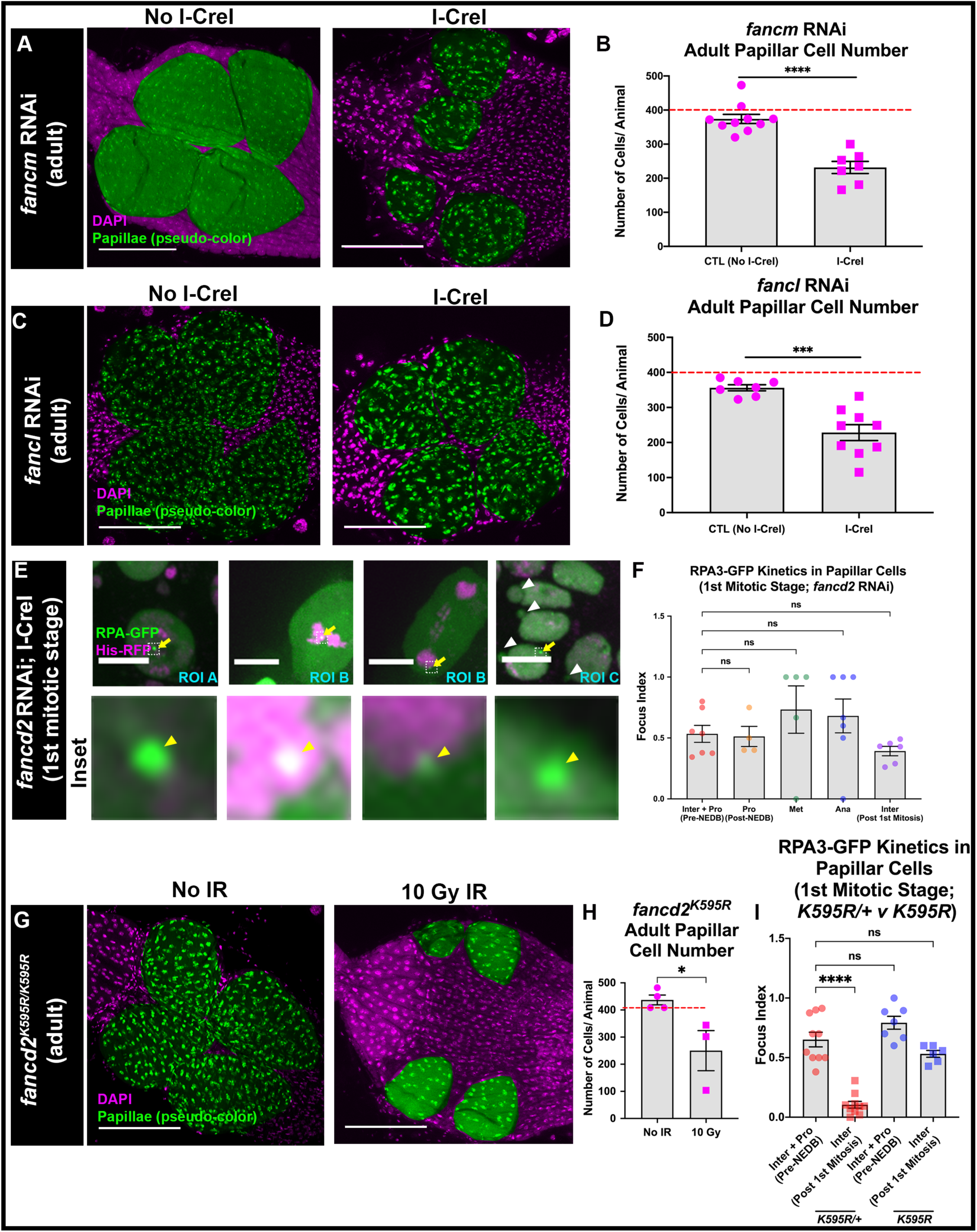
FA proteins are required for proper RPA3+ foci dynamics and cell survival following DSBs. (**A**) Adult rectums of *fancm* RNAi animals +/− *hs-I-CreI*. Green= pseudo-colored rectal papillae, magenta= DAPI (DNA). Scale bars = 100 μm. (**B**) Quantification of adult papillar cell number in *fancm* RNAi animals +/− *hs-I-CreI*. Each data point represents one animal. Red dashed line indicates the expected number of papillar cells in a WT adult. Each condition has at least 2 replicates. Statistical test: Unpaired t-test, p<0.0001. (**C**) Adult rectums of *fancl* RNAi animals +/− *hs-I-CreI*. Green= pseudo-colored rectal papillae, magenta= DAPI (DNA). Scale bars = 100 μm (**D**) Quantification of adult papillar cell number in *fancl* RNAi animals +/− *hs-I-CreI*. Each data point represents one animal. Red dashed line indicates the expected number of papillar cells in a WT adult. Each condition has at least 2 replicates. Statistical test: Unpaired t-test, p=0.0003. (**E**) RPA3-GFP foci recruitment in *fancd2* RNAi expressing papillar cells +/− *hs*-*I-Cre*I. ROI=region of interest, letters indicate separate cells at a given timepoint for a single animal. RPA3+ foci are marked with yellow arrows. The hatched box highlights an area magnified 10X in the corresponding inset below each panel. Enlarged foci are marked with yellow arrowheads. Micronuclei are marked with white arrowheads. Scale bars = 10μm. (**F**) Quantification of RPA3-GFP foci recruitment in *fancd2* RNAi expressing papillar cells during mitosis +/− *hs*-*I-Cre*I. Ordinary one-way ANOVA, p-value=0.7076. (**G**) Adult rectums of *fancd2^K595R^* animals +/− *hs-I-CreI*. Green= pseudo-colored rectal papillae, magenta= DAPI (DNA). Scale bars = 100 μm (**H**) Quantification of adult papillar cell number in *fancd2^K595R^* animals +/− IR. Each data point represents one animal. Red dashed line indicates the expected number of papillar cells in a WT adult. Each condition has at least 2 replicates. Statistical test: Unpaired t-test, p=0.0355. (**I**) Quantification of RPA3-GFP foci recruitment in *fancd2^K595R/+^* and *fancd2^K595R^* papillar cells before and after the first mitotic division +/− *hs*-*I-Cre*I. Ordinary one-way ANOVA, p<0.0001. See Methods for statistical notations.

